# Hypoxia is a dominant remodeler of the CD8^+^ T cell surface proteome relative to activation and regulatory T cell-mediated suppression

**DOI:** 10.1101/2021.06.07.447379

**Authors:** James R. Byrnes, Amy M. Weeks, Eric Shifrut, Julia Carnevale, Lisa Kirkemo, Alan Ashworth, Alexander Marson, James A. Wells

**Affiliations:** Department of Pharmaceutical Chemistry, University of California, San Francisco, San Francisco, CA, USA; Department of Biochemistry, University of Wisconsin-Madison, Madison, WI, USA; Department of Microbiology and Immunology, University of California, San Francisco, San Francisco, CA, USA; Gladstone Institutes, San Francisco, CA, USA; Department of Medicine, University of California, San Francisco, San Francisco, CA, USA; Innovative Genomics Institute, University of California, Berkeley, Berkeley, CA, USA; Parker Institute for Cancer Immunotherapy, San Francisco, CA, USA; Chan Zuckerberg Biohub, San Francisco, CA, USA; The Helen Diller Family Comprehensive Cancer Center, University of California, San Francisco, San Francisco, CA, USA; Department of Cellular and Molecular Pharmacology, University of California, San Francisco, San Francisco, CA, USA

## Abstract

Immunosuppressive factors in the tumor microenvironment (TME) impair T cell function and limit the anti-tumor immune response. T cell surface receptors that influence interactions and function in the TME are already proven targets for cancer immunotherapy. However, surface proteome remodeling of primary human T cells in response to suppressive forces in the TME has never been characterized systematically. Using a reductionist cell culture approach with primary human T cells and SILAC-based quantitative cell surface capture glycoproteomics, we examined how two immunosuppressive TME factors, regulatory T cells (Tregs) and hypoxia, globally affect the activated CD8^+^ surface proteome (surfaceome). Surprisingly, the CD8^+^/Treg co-culture only modestly affected the CD8^+^ surfaceome, but did reverse several activation-induced surfaceomic changes. In contrast, hypoxia dramatically altered the CD8^+^ surfaceome in a manner consistent with both metabolic reprogramming and induction of an immunosuppressed state. The CD4^+^ T cell surfaceome similarly responded to hypoxia, revealing a novel hypoxia-induced surface receptor program. Our findings are consistent with the premise that hypoxic environments create a metabolic challenge for T cell activation, which may underlie the difficulty encountered in treating solid tumors with immunotherapies. Together, the data presented here provide insight into how suppressive TME factors remodel the T cell surfaceome and represent a valuable resource to inform future therapeutic efforts to enhance T cell function in the TME.

## INTRODUCTION

Cytotoxic CD8^+^ T cells promote tumor cell killing, a function that is substantially modulated in the tumor microenvironment (TME)^1^. The TME is a complex mixture of tumor, somatic, and immune cells that create a unique environment in and around the tumor. Tumor-infiltrating CD8^+^ T cells encounter many cell types in the TME, including immunosuppressive regulatory T cells (Tregs).^2,3^ Tregs express immune suppressors such as CTLA-4^4^ and produce immunosuppressive adenosine^5^ and cytokines such as TGFβ, interleukin (IL)-10^6^, and IL-35^7^. Furthermore, Tregs and CD8^+^ cells compete for IL-2 for proliferation; high expression of the high affinity IL-2 receptor on Tregs allows them to out-compete CD8^+^ T cells for available IL-2.^8^ Tregs can also directly kill CD8^+^ T cells via the perforin pathway.^9^ Consistent with these many immunosuppressive activities, increased Treg tumor infiltration is associated with poor prognosis in a number of cancers, including non-small cell lung, hepatocellular, renal cell, breast, cervical, ovarian, and gastric cancers, as well as melanoma.^3,10,11^

Another hallmark of the TME is hypoxia due to poor and variable vascularization within the tumor.^1,12^ Hypoxia is common in the core of tumors and induces dramatic transcriptional changes.^1,12–17^ Tumor-associated hypoxia strongly influences the function of numerous immune cells from both the myeloid^18^ and lymphoid lineages^19–21^, with both stimulatory and inhibitory effects reported^22^. Hypoxia induces expression of CD39 and CD73 that catalyze the production of immunosuppressive adenosine from ATP.^23^ Hypoxia also induces the Warburg effect, which leads to tumor acidification, decreased CD8^+^ T cell proliferation, and reduced cytotoxic activity.^24^ Hypoxia additionally promotes recruitment of Tregs to the tumor^24^, and CD8^+^ cells have been observed to be excluded from areas of tumor hypoxia^25^. Recently, hypoxia was also linked to T cell exhaustion^26^. Inhospitable, hypoxic regions in solid tumors may also limit the function of CAR-T cells^27^, which could contribute to the limited success of targeting solid tumors with CAR-T cells.

The cell surface proteome, or surfaceome, mediates T cell interactions with the external environment, and the effect of external environmental factors on the T cell surfaceome has not yet been studied globally. Not only does the surfaceome help T cells sense and respond to the environmental conditions of the TME, but membrane proteins are useful surface markers and key regulators of the anti-tumor function of CD8^+^ cells. For example, proteins such as PD1 play crucial roles in the suppression of CD8^+^ cells.^28^ Consequently, many current immunotherapies target and modulate T cells through blockade or engagement of surface proteins (e.g. anti-PD1 or anti-CTLA-4 therapy, bispecific T cell engagers).^29^ Therefore, profiling how the CD8^+^ surfaceome changes in response to TME factors, such as Treg-mediated suppression or hypoxia, should expand our understanding of the basic biological response to these modulators.

We have taken a reductionist cell culture approach to begin to understand how Tregs and hypoxia modulate the cell surface proteome of primary CD8^+^ T cells. We first identified global and bi-directional changes in the CD8^+^ T cell surfaceome following classic activation with agonistic antibodies to CD3 and CD28 using quantitative cell surface capture mass spectrometry methods^30,31^. We discovered that co-culturing with Tregs, and especially hypoxic culture, significantly alter the activated CD8^+^ surfaceome in a manner consistent with reduced CD8^+^ activation. Although our *in vitro* model conditions are much less complex than what would be found in an *in vivo* TME, this approach allowed us to control and separately assess the impact of T cell activation, Treg co-culture, and hypoxia on the CD8^+^ T cell surfaceome. Collectively, these findings help illuminate how surfaceomic remodeling contributes to suppression of CD8^+^ T cells in the presence of Tregs and hypoxia, and provides a resource to identify potential markers for selective therapeutic targeting of suppressed CD8^+^ T cells in the TME or the design of new cellular therapies to overcome TME-mediated suppression.

## RESULTS

### Activation dramatically alters the CD8^+^ T cell surfaceome in a bi-directional fashion

Our strategy to study how the CD8^+^ cell surfaceome responds to activation, and how this response is altered by Treg suppression or hypoxia, is shown in Figure 1. Primary CD8^+^ cells were isolated from healthy donors and expanded 10- to 100-fold with anti-CD3 and anti-CD28 stimulation in the presence of IL-2. Cells were grown in medium containing light or heavy isotope-labeled lysine and arginine to quantitatively compare stimulation conditions using SILAC (stable isotopic labeling with amino acids in cell culture) coupled with a glycoprotein cell surface capture technique and LC-MS/MS^30–32^ (Figure 1A). This led to quantitative and uniform labeling as assessed by isotope distribution on four abundant and constitutively-expressed proteins (Supplemental Figure 1). We first compared the activation-induced changes in the CD8^+^ surfaceome before and after activation with anti-CD3/anti-CD28 for three days, and then examined how the program was altered by the addition of primary Tregs or hypoxic culture (Figure 1B).

**Figure 1.**
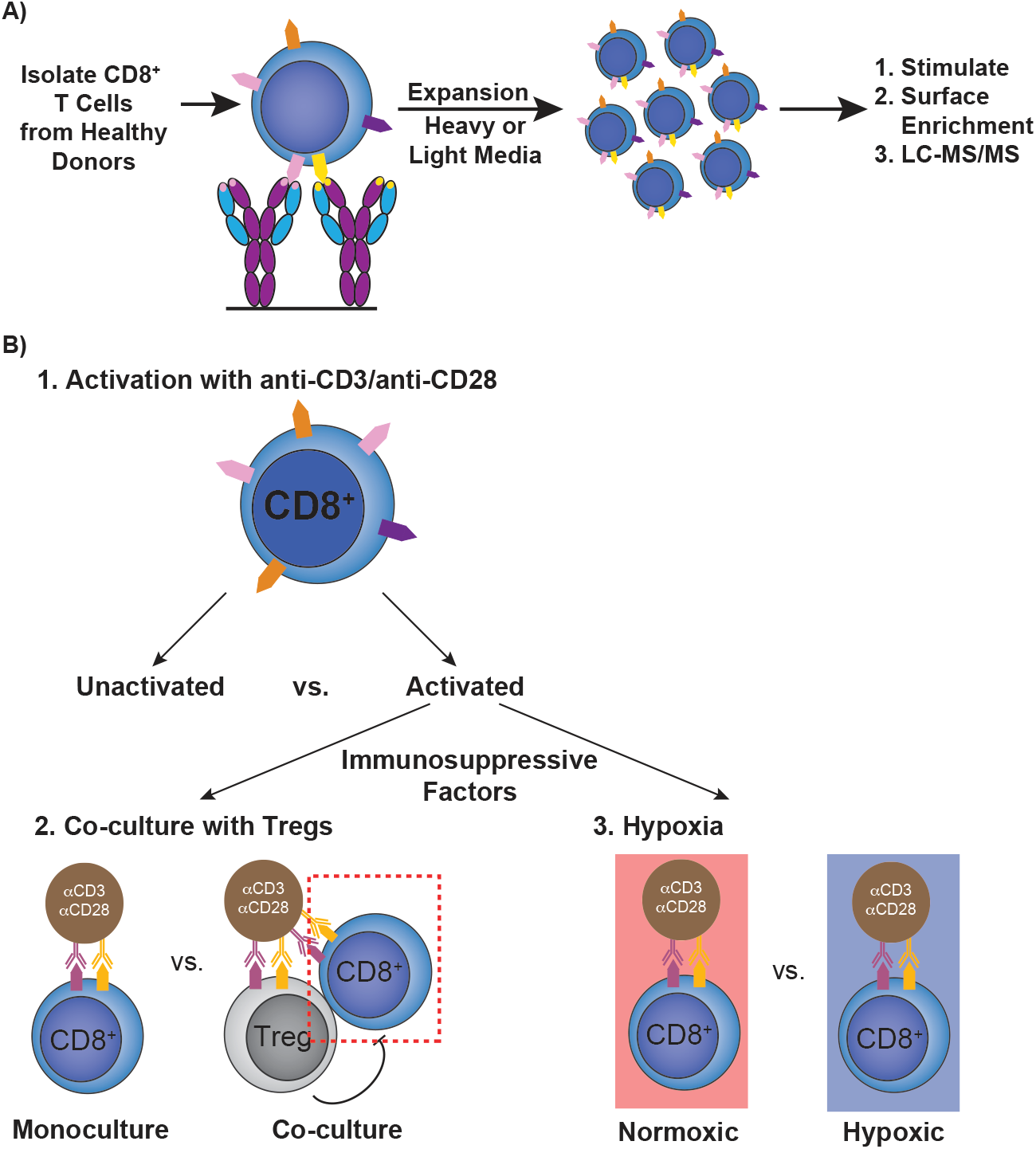
Overall strategy for cell surface glycoproteomic characterization of primary CD8^+^ T cells under various conditions. (A) Schematic depicting expansion and SILAC labeling workflow. Primary human CD8^+^ T cells were isolated and expanded using anti-CD3/anti-CD28 stimulation in media supplemented with IL-2 and either heavy or light arginine and lysine. After expansion, cells were stimulated in varying conditions before surface protein enrichment and protein identification with LC-MS/MS. (B) Strategy for assessing the effect of immunosuppressive stimuli on the activated CD8^+^ T-cell surfaceome. First, the surfaceomic changes associated with CD8^+^ activation in monoculture under normoxic conditions were analyzed. These changes then served as a baseline for later experiments examining the surfaceomic consequences of activating CD8^+^ T cells in co-culture with primary Tregs or in hypoxic culture.

Surfaceomic analysis of unstimulated and anti-CD3/anti-CD28-stimulated CD8^+^ T cells from four donors identified a total of 669 surface proteins (Figure 2A, Supplemental Table 1). Although there was donor-to-donor variation (Supplemental Figure 2), assessment of a compiled dataset including fold-change data from the four donors revealed about 16% of these proteins (106/669) consistently showed significant (*P*<0.05) 1.5-fold up- or downregulation. These significantly-altered proteins showed strong correlation between most donors (Supplemental Figure 2). We observed changes in classic markers of T-cell activation, including upregulation of two classic T cell activation markers (CD69^33^ and the transferrin receptor [TFRC]^34^), and downregulation of the IL-7 receptor (internalized upon activation) and VIPR1^35^ (Figure 2B).

**Figure 2.**
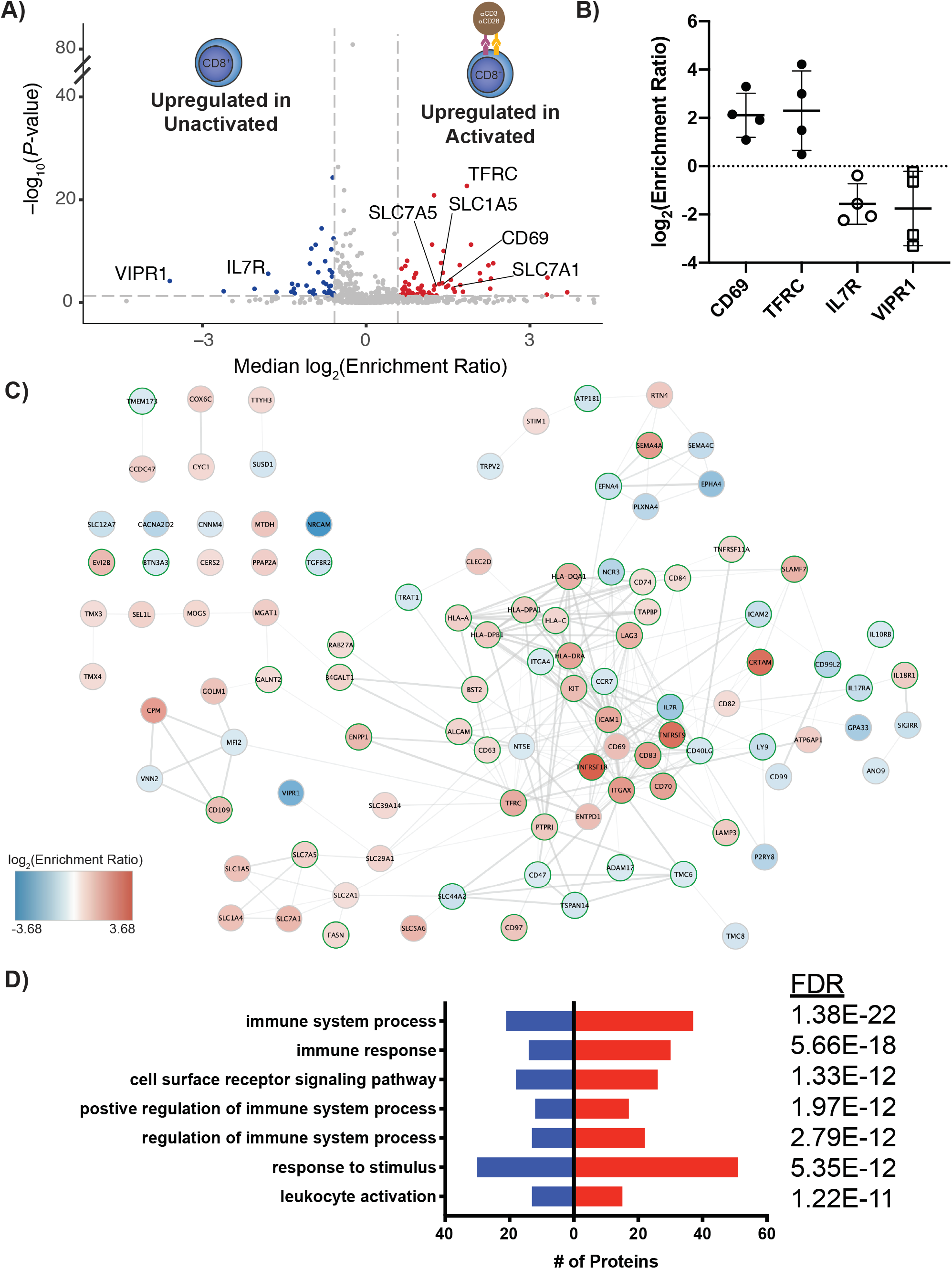
Surface proteomics reveals both well-established and novel activation-induced changes in surface protein levels. (A) Volcano plot of surface protein changes following stimulation of CD8^+^ T cells with anti-CD3/anti-CD28 beads. Data represent compiled results from N=4 donors. Proteins with a −/+1.5-fold change and P<0.05 were included in downstream analysis. Proteins significantly down-(blue) or upregulated (red) are indicated. (B) Log_2_(Enrichment Ratio) of indicated proteins. Each dot represents data from an individual donor. Line represents the mean and error bars are standard deviation. (C) STRING analysis of all significantly-altered proteins in (A). Network is overlaid with a color gradient representing log_2_(Enrichment Ratio) for each individual protein. Proteins with a gene ontology (GO) biological process annotation of “immune system process” are indicated with green borders. (D) Significantly-altered proteins were subjected to GO biological process pathway analysis using the STRING database. The number of proteins identified, the direction of regulation, and analysis FDR for each process are indicated.

Large-scale network analysis of the significantly-altered proteins demonstrates the roughly symmetrical, bi-directional response of the T cell surfaceome to activation, with 66 proteins upregulated and 40 proteins downregulated (Figure 2C). In addition to CD69 and TFRC, numerous well-established T cell activation markers were also upregulated, including CD63^33^, CD83^33^, CD97^36^, and CD109^37^. Importantly, multiple solute carrier (SLC) transporters were also upregulated on activated CD8^+^ T cells, including the amino acid transporters SLC1A5 and SLC7A5 that have been previously implicated in supporting T cell activation.^38^ Of note, comparison of our proteomics data with RNAseq data from activated versus resting CD8^+^ cells in the DICE database revealed a loosely positive correlation (R=0.25, *P*<0.0001, Supplemental Figure 3A).^39^ It is well known that protein and RNA levels show only mild correlations because of differences in stability and regulation. Assessment of only proteins that were significantly changed in our proteomics data revealed a stronger correlation (R=0.53, *P*<0.0001, Supplemental Figure 3B). However, several proteins that demonstrated significant change in our data showed minimal or divergent change in the RNAseq data (e.g. CD70, 2.09 vs. −0.41; SLC5A6, 1.62 vs. 0.46; IL7R, −1.80 vs. 0.73 [log_2_(enrichment ratio) for proteomics vs. RNAseq, respectively]). These discrepancies could be due to differential activation conditions, but nonetheless underscore the importance of protein-level profiling to capture surface protein remodeling in immune cells.

More globally, pathway analysis of up- and downregulated cell surface proteins revealed the most significant enrichment for proteins implicated in immune function, with a slight trend towards upregulation of these proteins (Figure 2D, proteins annotated for GO.0002376: immune system process are indicated with green borders in Figure 2C). Collectively, these surfaceomic data identify classic (e.g. CD69) and some newly-recognized (e.g. integrin αX [ITGAX], SLC39A14, BST2) markers for immune activation of primary CD8^+^ T cells.

### Co-culture with Tregs modulates the activated CD8^+^ surfaceome

We next analyzed the effect that primary Tregs have upon the surfaceome of activated primary CD8^+^ T cells in a 1:1 ratio co-culture after three days (Figure 3A, Supplemental Table 2). Relative to activation alone, Treg co-culture had a mild global impact on the surfaceome of activated CD8^+^ cells, with significant (*P*<0.05) up- or downregulation (−/+ 1.5-fold change) of only 34 out of 675 proteins detected (Figure 3A). Changes in these proteins were again largely consistent between donors (Supplemental Figure 4). Of note, upregulation of TFRC and the pro-inflammatory cytokine receptor IL18R1 on the cell surface was blunted by the addition of Tregs (Figure 3B). In the presence of Tregs, IL7R showed variable upregulation between donors, opposite of the trend seen with cell activation in monoculture (Figures 2B and 3B). Similarly, L-selectin (SELL), a protein which is typically downregulated with T cell activation^40^, was slightly upregulated on CD8^+^ T cells, again consistent with a suppressive effect of Tregs (Figure 3B). Due to the smaller number of significantly changed proteins in this dataset, network analysis was not as striking as in the activation dataset (Figure 3C). However, in contrast to the monoculture CD8^+^ T cell activation dataset, which demonstrated an upward trend in proteins implicated in immune processes, there is a downward trend in immune-annotated proteins when CD8^+^ cells are activated in the presence of Tregs, consistent with an immunosuppressive effect (Figure 3D). Furthermore, many proteins that were upregulated on activated CD8^+^ T-cells in monoculture (Figure 2) were downregulated upon activation in the presence of Tregs (11 of 23 downregulated proteins, Figure 3C). Among these proteins are several SLCs, including SLC1A5 and SLC7A1. Collectively, these data suggest that the presence of Tregs reverses at least part, but not all, of the activation-induced surfaceomic response in CD8^+^ T cells.

**Figure 3.**
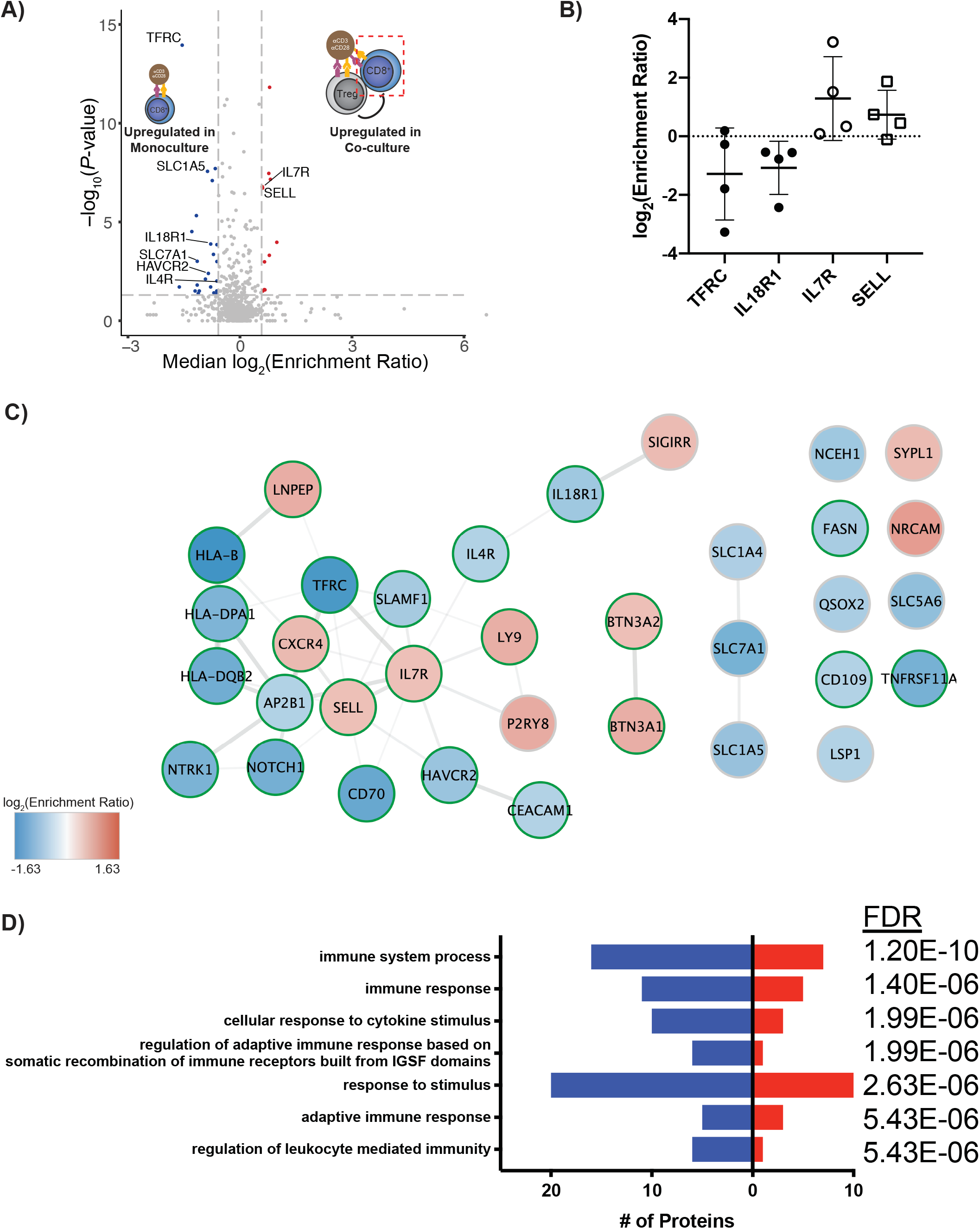
Treg co-culture causes significant changes in the surfaceome of activated CD8^+^ T cells consistent with immunosuppression. (A) CD8^+^ T cells were stimulated with anti-CD3/anti-CD28 beads either in the absence or presence of Tregs. Following culture, CD8^+^ cells were isolated for cell surface proteomics. Volcano plot shows compiled results from N=4 donors. Proteins with a −/+1.5-fold change and P<0.05 were included in downstream analysis. Proteins significantly down-(blue) or upregulated (red) are indicated. (B) Log_2_(Enrichment Ratio) of indicated proteins. Each dot represents data from an individual donor. Line represents mean and error bars are standard deviation. (C) STRING analysis of all significantly-altered proteins in (A). Network is overlaid with a color gradient representing log_2_(Enrichment Ratio) for each individual protein. Proteins with a gene ontology (GO) biological process annotation of “immune system process” are indicated with green borders. (D) Significantly-altered proteins were subjected to GO biological process pathway analysis using the STRING database. The number of proteins identified, the direction of regulation, and analysis FDR for each process are indicated.

### Hypoxia triggers large-scale surfaceomic changes in activated CD8^+^ T cells consistent with immunosuppression and anaerobic reprogramming

We next wanted to test how hypoxia affects the surfaceome of activated CD8^+^ T cells. Interestingly, over three days in culture, activated CD8^+^ T cells proliferated only ~1.3-fold faster in normoxia (20% O_2_) compared to hypoxia (1% O_2_), and there were no substantial differences in cell viabilities (Supplemental Figure 5). However, surface proteomics of CD8^+^ T cells activated in normoxic or hypoxic conditions revealed substantial remodeling of the surface proteome. Of a total of 1064 proteins identified, 196 were significantly (*P*<0.05) up- or down-regulated (−/+ 1.5-fold change, Figure 4A, Supplemental Table 3) in hypoxia relative to normoxia. The fold changes observed for these significantly-altered proteins showed higher correlation among donors (Supplemental Figure 6) than seen in the activation (Supplemental Figure 2) and Treg co-culture (Supplemental Figure 4) datasets. The upregulation of the hypoxia-induced glucose transporter SLC2A3 (GLUT3)^41^ we observed is consistent with a shift towards glycolysis. We also observed downregulation of activin receptor type-1 (ACVR1), which is sequestered in endosomes under hypoxic conditions^42^ (Figure 4B).

**Figure 4.**
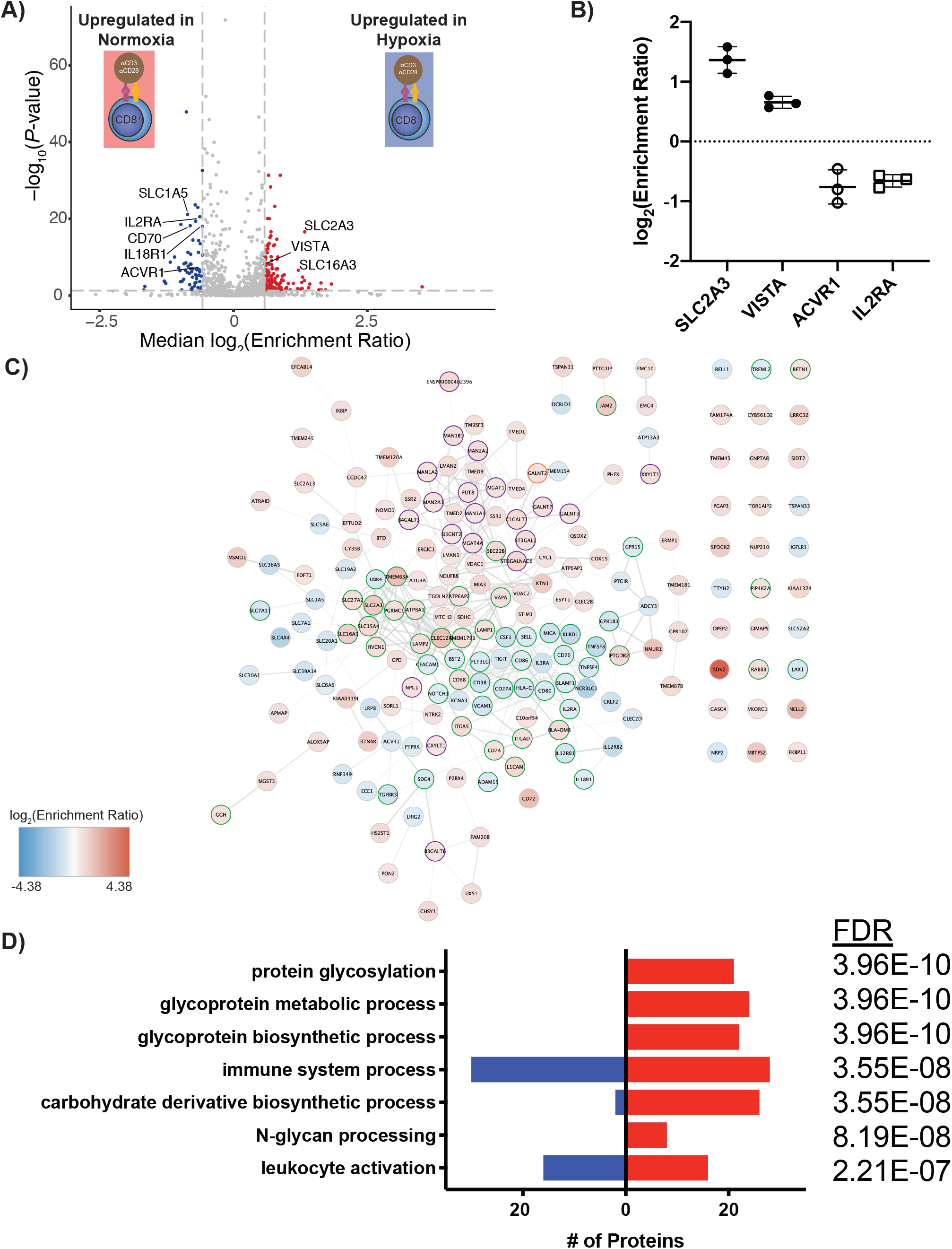
Hypoxia induces surfaceomic changes representative of both immunosuppression and a general response to hypoxia. (A) CD8^+^ T cells were stimulated with anti-CD3/anti-CD28 beads either normoxic (20% O_2_) or hypoxic (1% O_2_) for three days. Volcano plot shows compiled results from N=3 donors. Proteins with a −/+1.5-fold change and P<0.05 were included in downstream analysis. Proteins significantly down-(blue) or upregulated (red) are indicated. (B) Log_2_(Enrichment Ratio) of indicated proteins. Each dot represents data from an individual donor. Line represents mean and error bars are standard deviation. (C) STRING analysis of all significantly-altered proteins in (A). Network is overlaid with a color gradient representing log_2_(Enrichment Ratio) for each individual protein. Proteins with a gene ontology (GO) biological process annotation of “immune system process” are indicated with green borders and proteins with an annotation of “protein glycosylation” with purple borders. Proteins annotated for both processes are indicated with an orange border. (D) Significantly-altered proteins were subjected to GO biological process pathway analysis using the STRING database. The number of proteins identified, the direction of regulation, and analysis FDR for each process are indicated.

Network visualization of proteins significantly altered by hypoxia revealed multiple clusters of up- and down-regulated proteins (Figure 4C). Interestingly, gene ontology analysis did not identify significant enrichment for proteins involved in response to hypoxia (GO.0001666) or cellular response to hypoxia (GO.0071456). However, four of these significantly-altered proteins are found in the “Hallmark Hypoxia” gene set (SDC4, SLC6A6, B3GALT6, and SLC2A3).^43^ We did observe marked upregulation of proteins involved in protein glycosylation and glycoprotein metabolic processes (Figure 4C and D). Tumor hypoxia is well established to cause glycan remodeling of tumor cell surface proteins^44^, and our data suggest the same may be true for hypoxic T cells. Among the 132 hypoxia-upregulated proteins were numerous proteins involved in the unfolded protein response (e.g. ERMP1^45^, FKBP11^46^, MBTPS2^47^) and regulation of autophagy (ATG9A^48^, ERGIC1, SEC22B^49^). In contrast, proteins implicated in immune function were strongly represented in the pool of 64 hypoxia-suppressed proteins (Figure 4C and D). In addition to the late activation marker IL2RA (CD25), co-stimulatory receptors, such as CD80 and CD86, as well as receptors for interleukins-3, -12, and -18 were significantly downregulated in hypoxic conditions (Figure 4B and C). Several SLCs implicated in T cell function, including SLC1A5^38^, and the pro-proliferative cytokines TNFSF4 (OX40 ligand) and TNFSF8 (CD30 ligand) were also downregulated. Of note, hypoxia led to downregulation of some immunosuppressive proteins, including CD70^50^ and TIGIT^51^, but consistent upregulation of the checkpoint molecule VISTA (C10orf54)^52^ (Figure 4B and C). To validate these observations, we performed flow cytometry for IL18R1 and CD70 on CD8^+^ cells from four different donors activated in normoxia and hypoxia. Both proteins were upregulated on normoxic activated cells relative to resting cells, although the degree of upregulation was variable (Supplemental Figure 7). Hypoxia resulted in blunted cell surface induction of both proteins (Supplemental Figure 7), consistent with our proteomics results. Collectively, these data reveal that hypoxia induces dramatic remodeling of the activated CD8^+^ T cell surfaceome and markedly regulates numerous proteins important for T cell activation.

### CD8^+^ and CD4^+^ T cells demonstrate similar hypoxia-induced surface remodeling

Given the dramatic effect of hypoxic culture on the surface proteome of activated CD8^+^ T cells, we next determined how hypoxia modulates surface protein expression on another cell subset important for the anti-tumor immune response: CD4^+^CD25^−^ conventional T cells. Analysis of CD4^+^CD25^−^ T cells from the same donors as in Figure 4 again revealed dramatic surfaceomic remodeling when these cells were activated in hypoxic conditions (Figure 5A, Supplemental Table 4). Of the 1144 proteins identified, 992 were also identified in our CD8^+^ hypoxia dataset. The fold-change ratios of these commonly-identified proteins showed significant correlation (R=0.82, *P*<0.0001) between CD8^+^ and CD4^+^CD25^−^ cells (Figure 5B). Interestingly, more proteins were significantly up-(306) or downregulated (204) in the CD4^+^CD25^−^ dataset, and the magnitude of these changes was larger than those observed for the CD8^+^ cells. However, there was a large overlap in the sets of significantly-altered proteins (Figure 5C and D). Consequently, functional enrichment for these overlapping proteins was similar to that observed when analyzing the CD8^+^ response alone, with significant downregulation of proteins involved in immune-related processes (e.g. CD70 and IL18R1) and upregulation of proteins involved in solute transport (e.g. SLC2A3, SLC16A3) and protein glycosylation. Taken together, these data show there is substantial similarity in the surfaceomic response of CD8^+^ and CD4^+^ T cells to hypoxia.

**Figure 5.**
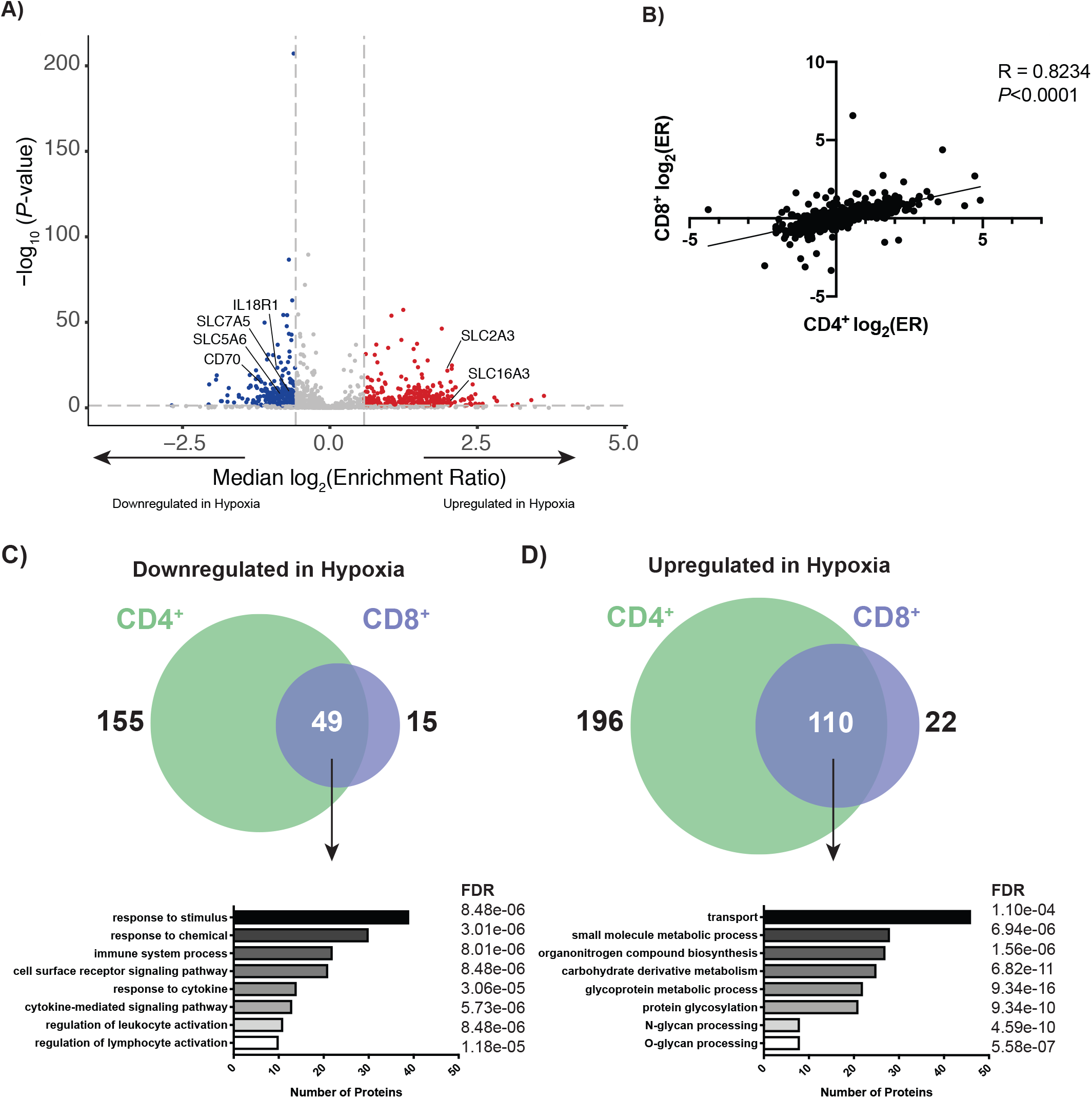
CD8^+^ and CD4^+^ surface proteomes respond similarly to hypoxia. (A) CD4^+^CD25^−^ T cells were stimulated with anti-CD3/anti-CD28 beads either normoxic (20% O_2_) or hypoxic (1% O_2_) for three days. Volcano plot shows compiled results from N=3 donors. Proteins with a −/+1.5-fold change and P<0.05 were included in downstream analysis. Proteins significantly down-(blue) or upregulated (red) are indicated. (B) Spearman correlation comparing the log_2_(Enrichment Ratio [ER]) for both CD8^+^ and CD4^+^ T cells activated under hypoxic conditions. Venn diagrams showing proteins commonly down-(C) or upregulated (D) in hypoxia on both CD8^+^ and CD4^+^ cells. Below each Venn diagram is are results from a GO biological process pathway analysis for commonly regulated proteins using the STRING database. The number of proteins identified and analysis FDR for each process are indicated.

### Analysis of all CD8^+^ datasets reveals a conserved response to immunosuppressive stimuli

We next cross-referenced the observed surfaceomic changes associated with Treg co-culture and hypoxia. Interestingly, gene set enrichment analysis comparing the expression level of all proteins identified in our datasets revealed significant enrichment for a number of biological processes, namely organic and amino acid transmembrane transport (Figure 6A). Consistent with this observation, analysis of proteins significantly upregulated with CD8^+^ activation but blunted due to hypoxia or Tregs identified three solute carriers (Figure 6B). Of these, SLC1A5 and SLC7A1 have been previously implicated in supporting T cell function following activation. Furthermore, the pro-inflammatory cytokine receptor IL18R1 and potentially inhibitory CD70^53^ that were upregulated with CD8^+^ activation exhibited blunted surface induction with hypoxia and Tregs. Strikingly, when examining proteins downregulated upon CD8^+^ activation in standard conditions but upregulated with hypoxia and Tregs, there are no commonly regulated proteins (Figure 6C). This suggests that protein upregulation seen during activation in hypoxic conditions may be primarily a general response to hypoxia. Together, the downregulation of a small but common set of surface proteins by both Treg co-culture and hypoxia may represent a conserved response of the T cell surface proteome to these two immunosuppressive stimuli.

**Figure 6.**
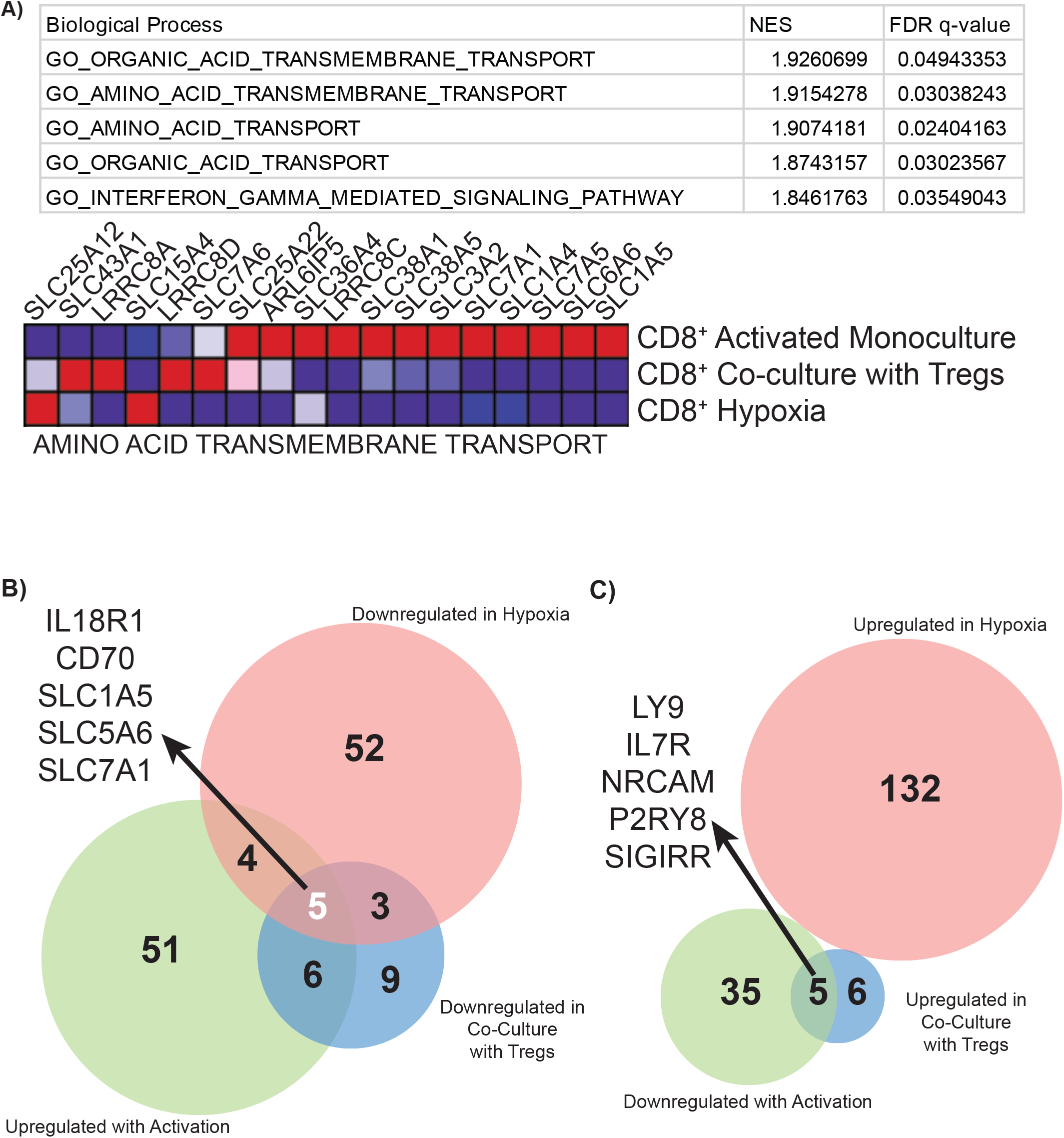
Analysis of all datasets reveals commonalities in the effects of Tregs and hypoxia on the activated CD8^+^ T cell surfaceome. (A) Gene set enrichment analysis (GSEA) output of all commonly-detected proteins in all three datasets. The five biological processes with FDR q-values <0.05 are shown, as are the normalized enrichment scores (NES) for each process. A heat map of proteins associated with GO_AMINO_ACID_TRANSMEMBRANE_TRANSPORT is also shown with the three datasets indicated. Red and blue indicate high and low enrichment, respectively. (B) Venn diagram showing intersection of proteins that were upregulated upon CD8^+^ activation, but downregulated by Treg co-culture or hypoxia. The five proteins at the intersection of all three datasets are indicated. (C) Venn diagram showing intersection of proteins downregulated upon CD8^+^ activation, but upregulated by Treg co-culture or hypoxia.

## DISCUSSION

A major obstacle encountered by CD8^+^ T cells when mounting an anti-tumor immune response is the immunosuppressive TME. Although Tregs and hypoxia are known to be immunosuppressive, we have little global understanding of how these affect the CD8^+^ T cell surface proteome. Our studies illuminate the bi-directional surfaceomic changes on CD8^+^ T cells associated with cell activation and with immunosuppression following co-culture with Tregs or hypoxic culture. The presence of Tregs partly reversed activation-induced changes in CD8^+^ T cells, consistent with suppressed cell activation. Perhaps surprisingly, the effect of hypoxia was much larger. Hypoxia triggered large-scale surfaceomic remodeling consistent with a general cellular response to the new metabolic demands of a low oxygen environment. Most importantly, cross-referencing of the effects of both Treg co-culture and hypoxia on the surface proteome of activated CD8^+^ T cells exposed a small, but intriguing list of common changes in the expression of proteins primarily involved in nutrient transport. This finding suggests that induced downregulation of surface proteins important to support CD8^+^ activation may be one mechanism of immunosuppression in the TME.

Global cell-surface capture proteomics not only provided a means of visualizing upregulation of known activation markers to validate our experimental system, but also generated an unbiased list of additional proteins demonstrating activation-associated changes. Importantly, our approach identified significant cell surface upregulation of numerous well-established activation markers (e.g. CD69, TFRC, CRTAM) that support T cell function. We also observed upregulation of TNFRSF18 (GITR), a molecule vital for CD8^+^ anti-tumor function and agonism of which synergizes with anti-PD1 checkpoint therapy.^54^ The exhaustion marker LAG3 was upregulated, as was ENTPD1 (CD39), which has been reported to increase following T cell activation^55^. Upregulation of CD39 is especially interesting because it facilitates conversion of extracellular ATP to ADP, which is the first step to generating immunosuppressive adenosine. Although this could represent a negative feedback loop to dampen T cell activation, we observed that CD73 (NT5E), which converts ADP to adenosine, was downregulated following activation. This is consistent with the recent observation that following activation human T cells strongly upregulate CD39, whereas CD73 remains near baseline levels.^55^

Another class of proteins that demonstrated divergent trends are cell adhesion molecules. Many were significantly upregulated upon activation, including CD84^56^, ALCAM, which stabilizes the immunological synapse^57^, and integrin αX, which is associated with improved migratory potential^58^. However, NRCAM, ICAM1, and ITGA4, the latter of which supports T cell migration^59^, were downregulated. The polarity of this response following activation may be representative of the complex interactions an activated T cell must make with its environment to not only infiltrate an immunologically-active zone, but also interact with target cells. Lastly, we observed significant upregulation of a number of solute transporters involved in transporting amino acids, vitamins, and other nutrients. Of these, several (SLC29A1, SLC2A1, SLC7A1) have previously been implicated in supporting T cell function, and are crucial to help T cells respond to the metabolic demands following activation.^60,61^ However, the choline transporter SLC44A2 and potassium-chloride cotransporter SLC12A7 were downregulated. Collectively, these data not only validate our approach and provide a broad picture of the surfaceomic remodeling following activation, but also serve as a benchmark for assessing the effects of hypoxia and Tregs on the activated CD8^+^ surfaceome.

Activation of CD8^+^ T cells in the presence of Tregs did not induce dramatic surfaceomic shifts relative to the hypoxia or activation datasets. However, co-culture showed several of the classic markers up-regulated in activation, TFRC, SLC1A5, and SLC7A1^38^, are downregulated upon addition of Tregs. Of note, the exhaustion marker HAVRC2 (TIM-3) was also downregulated on CD8^+^ T cells activated in the presence of Tregs. This may be a consequence of diminished CD8^+^ activation in the presence of Tregs. As our experiments were limited to three days of co-culture, the trends we observe likely represent changes that occur relatively early in an environment containing both CD8^+^ cells and Tregs. Longer-term culture conditions may demonstrate even more dramatic changes associated with prolonged Treg-mediated suppression of CD8^+^ cells.

Multiple studies have specifically examined the direct effect of hypoxia on CD8^+^ T cells, either by modulating the activity of the canonical hypoxia-associated transcription factor hypoxia-inducible factor-1α (HIF-1α) or using hypoxic cell culture. Doedens et al. showed that knockdown of the negative HIF regulator VHF enhanced the cytotoxic signature of CD8^+^ cells and led to sustained effector function.^62^ Similarly, Gropper et al. showed that CD8^+^ T cells cultured in 1% O_2_ exhibited enhanced cytolytic activity.^63^ However, these cells proliferated only half as quickly as normoxic cells during the culture period, which may counteract the enhanced effector function of these cells. Taken with the results of other studies examining CD8^+^ function in hypoxia, low oxygen tension appears to exert a net immunosuppressive effect on the antitumor response.^22^ Our data add to these focused functional studies and revealed that hypoxia led to the most substantial surfaceomic changes of all conditions tested with respect to the number of proteins demonstrating significant change. Additionally, the response of CD4^+^ T cells to hypoxia was tightly correlated with that of CD8^+^ cells, suggesting hypoxia has similar impacts on both cell types.

Analysis of the altered surface proteins in hypoxic culture (e.g. GLUT3 upregulation), reveals changes consistent with a metabolic change toward glycolysis. One protein that showed robust upregulation in both CD8^+^ and CD4^+^CD25^−^ cells was SLC16A3, a hypoxia-induced lactate transporter that helps export lactate produced from glycolysis which has previously been implicated in supporting tumor growth.^64^ Interestingly, the related SLC16A1 was recently shown to help intratumoral Tregs metabolically cope with high lactate levels in areas of tumor hypoxia.^65^ The role of SLC16A3 upregulation on hypoxic CD8^+^ and CD4^+^CD25^−^ cells remains unclear and further functional follow up is needed, but previous studies indicate inhibition of this transporter can enhance effector function.^66^ Intratumoral hypoxia is also known to dramatically alter the glycoproteome of tumor cells, changes that in turn regulate cell migration and metastasis. Our data suggest the same could be true for T cells, which may have similar consequences for T cell migration within the tumor and could represent another feature that could be harnessed to selectively target hypoxic T cells. Further studies are needed to profile the hypoxic T cell glycoproteome.

Intriguingly, proteins involved in hypoxia-induced autophagy (ATG9A^48^, ERGIC1, SEC22B^49^) and the unfolded protein response (ERMP1^45^, FKBP11^46^, MBTPS2^47^) were also upregulated. These two pathways are associated with a survival response under hypoxic conditions.^67^ However, activation of the unfolded protein response pathway is usually observed only at very low levels of oxygen (<0.1%)^67^, and is often associated with cell death. Although we did not observe rampant cell death in our experiments (performed at 1% oxygen), even lower oxygen levels within dividing T cell clusters or the compounded metabolic demands of hypoxic conditions and activation may have triggered these pathways. Interestingly, T cell activation has also been associated with induction of the endoplasmic reticulum stress response and autophagy.^68,69^ Collectively, these observations suggest that hypoxia places additional stress on the activation-associated metabolic shift, ER stress, and autophagic responses that are typically induced in T cells following activation. This added stress may impair T cell function and the ability for T cells to cope with sustained stimulation in the hypoxic TME.

Finally, one global observation from our hypoxic CD8^+^ data is that although a large number of proteins show significant change, the magnitude of these changes is mild, with only 28/196 (14%) of significantly-altered proteins showing a change of greater −/+ two-fold change. This may suggest that the T cell response to hypoxia results in a distributed biology, where many small changes in protein abundance collectively lead to a suppressed state. This complicates functional follow-up of proteins identified in this study, but nonetheless this will be a focus of future efforts to provide greater resolution of the surface protein changes that lead to hypoxia-induced T cell suppression.

Taken together, our data provide the opportunity to search for common surfaceomic changes in response to both Treg co-culture and hypoxia. Gene set enrichment analysis comparing the three datasets revealed very strong enrichment for transmembrane amino acid transport in the CD8^+^ monoculture activation dataset. These transporters are vital for fueling macromolecule production needed for both proliferation and effector function, with both the glutamate transporter SLC1A5 and amino acid transporter SLC7A1 previously implicated in supporting T cell activation. Interestingly, SLC5A6 is a sodium-dependent multivitamin transporter that has not been previously implicated in T cell function. Together, these three transporters may represent a common set of proteins whose expression is modulated by immunosuppressive factors. Downregulation of these transporters by the suppressive forces tested here suggests these factors blunt the T cell activation response by limiting the increased metabolism required to sustain proliferation and effector function (Figure 7). This premise is consistent with a recent study finding that hypoxia-induced metabolic stress promotes T cell exhaustion.^26^ It remains to be seen if these solute transporters are also downregulated on infiltrating T cells *in vivo*, but our findings begin to illuminate how TME-associated immunosuppression of CD8^+^ cells may result from effectively starving T cells as they attempt to mount an anti-tumor response.

**Figure 7.**
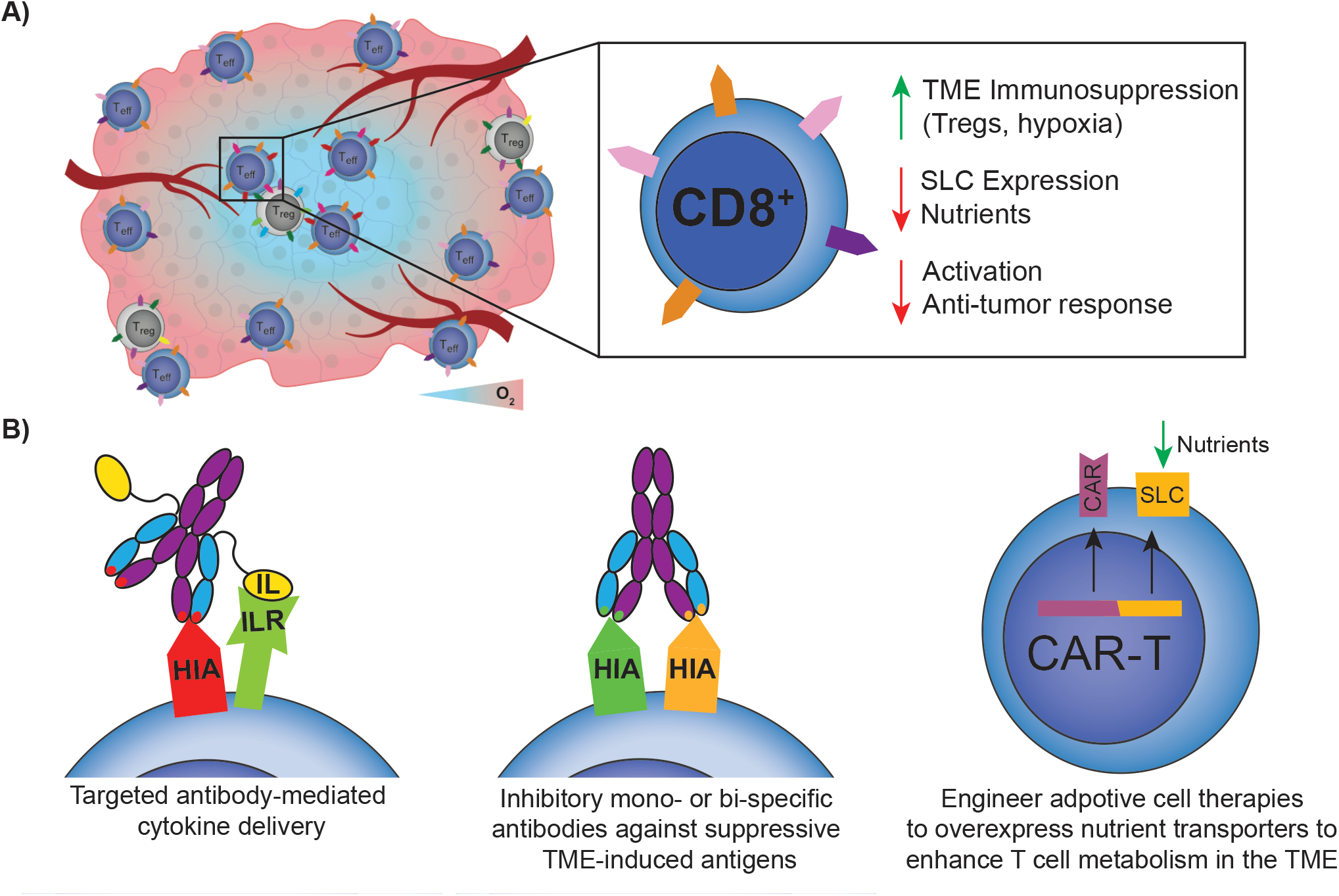
Metabolic model of CD8^+^ immunosuppression and potential therapeutic strategies. (A) Tregs and hypoxia suppress expression of nutrient transporters on CD8^+^ T cells, limiting T cell activation and the anti-tumor response. (B) Proteins identified in our studies could be utilized for targeted antibody-based therapies to enhance CD8^+^ function in the TME. Hypoxia-induced antigen (HIA) used as an example. Antibody-cytokine fusion targeting an HIA could be used to deliver pro-proliferative cytokines to CD8^+^ T cells (left). Alternatively, TME-induced antigens that inhibit T cell function could be blocked with mono- or bi-specific antibodies. Lastly, adoptive cellular therapies could be reprogrammed to express higher levels of SLCs to help cope with the metabolic stresses found within the TME, which may enhance the efficacy of these therapies against solid tumors.

In conclusion, our cell surface proteomics data revealed upregulation of well-established activation markers following CD8^+^ stimulation, but also revealed many additional upregulated proteins that may augment T cell effector function and help T cells cope with the increased metabolic demands associated with rapid proliferation. Both immunosuppressive factors tested caused significant changes to the CD8^+^ surfaceome consistent with diminished T cell function. Interestingly, however, these factors converged on mediating downregulation of transporters crucial for linking the T cell activation response and the metabolic shift that follows stimulation, uncovering a potential metabolic mechanism for immunosuppression in the TME. Future work will aim to validate and screen these proteins for potential strategies to support the function of T cells in hypoxic environments. Such strategies could either block the function of inhibitory proteins upregulated in the TME, or could mediate selective delivery of beneficial cytokines via antibody-cytokine fusions (Figure 7B). Furthermore, our results provide a starting point to identify proteins that could be modulated in adoptive T cellular therapies (e.g. CAR-T) to enhance function of these cells in the solid TME (Figure 7B). Collectively, our findings provide important insight into the plasticity of the T cell surfaceome and lay the foundation for future efforts to not only further characterize, but also potentially therapeutically engage, T cell surface proteins within the TME.

## ACKNOWLEDGEMENTS

We thank Kevin Leung for valuable insight and discussions and the other members of the Wells Lab for their support.

J.A.W. is grateful for funding from the Harry and Dianna Hind Endowed Professorship in Pharmaceutical Sciences, NIH R35GM122451; J.A.W., A.A. and A.M. were supported on a common grant from the Parker Institute for Cancer Immunology. Postdoctoral Fellowship support included a National Institutes of Health National Cancer Institute F32 (5F32CA239417 to J.R.B.). A.M. holds a Career Award for Medical Scientists from the Burroughs Wellcome Fund, is an investigator at the Chan Zuckerberg Biohub and is a recipient of The Cancer Research Institute (CRI) Lloyd J. Old STAR grant. The Marson lab has received funds from the Innovative Genomics Institute (IGI), the Simons Foundation and the Parker Institute for Cancer Immunotherapy (PICI).

## AUTHOR CONTRIBUTIONS

J.R.B. and A.M.W. designed the research, performed experiments, and analyzed data. E.S. and J.C. performed experiments. L.K. provided valuable technical assistance and training. A.A., A.M., and J.A.W. conceived and supervised the research. J.R.B. and J.A.W. co-wrote the manuscript, and all authors provided edits and approved the final version.

## METHODS

### Cell isolation

Primary human T cells were isolated from leukoreduction chamber residuals following Trima Apheresis (Blood Centers of the Pacific, San Francisco, CA) using established protocols^70^. Briefly, peripheral blood mononuclear cells (PBMCs) were isolated using Ficoll separation in SepMate tubes (STEMCELL Technologies, Vancouver, Canada) in accordance with the manufacturer’s instructions. CD8^+^ T cells were isolated from PBMCs using either the EasySep™ Human CD8^+^ T cell Isolation Kit or the RosetteSep™ Human CD8^+^ T Cell Enrichment Cocktail (STEMCELL), following the manufacturer’s protocol. CD4^+^CD25^−^ conventional T cells and CD4^+^CD127^low^CD25^+^ Tregs were isolated from PBMCs with the EasySep™ Human CD4^+^CD127^low^CD25^+^ Regulatory T Cell Isolation Kit (STEMCELL). Isolated cell populations were analyzed for purity by flow cytometry on a Beckman Coulter CytoFlex flow cytometer using a panel of antibodies (anti-CD3 [UCHT1], anti-CD4 [OKT4], anti-CD8a [RPA-T8], anti-CD25 [M-A251], anti-CD45RA [HI100], and anti-CD127 [A019D5], all from BioLegend, San Diego, CA). Pilot experiments comparing FACS sorting and magnetic bead separation showed similar results with respect to cell purity, so magnetic bead separation was used.

### Cell culture/SILAC labeling

Following isolation, cells were adjusted to 1e6 cells/mL in RPMI 1640 Medium for SILAC (ThermoFisher) supplemented with 1000 U/mL Penicillin/Streptomycin (Gemini Bio-Products), 10 mM HEPES (UCSF Cell Culture Facility), 100 μM non-essential amino acids (Lonza), 1 mM sodium pyruvate (VWR), 55 μM 2-mercaptoethanol (Gibco), 10 mM N-acetyl-cysteine (Sigma), and 10% dialyzed FBS (Gemini). Media was also supplemented with either light L-[12C6,14N2] lysine/L-[12C6,14N4] arginine (Sigma) or heavy L-[13C6,15N2] lysine/L-[13C6,15N4] arginine (Cambridge Isotope Laboratories). CD8^+^ and CD4^+^ T effectors were stimulated by addition to tissue culture flasks coated with anti-CD3 (produced in-house, clone OKT3) and anti-CD28 ([D2Z4E], Cell Signaling) antibodies or with anti-CD3/anti-CD28 Dynabeads (Thermo) at a 1:1 bead:cell ratio in the presence of 50 U/mL recombinant human IL-2 (Thermo). Following two days of culture, the stimuli were removed and cells allowed to expand in heavy or light SILAC media with 50 U/mL IL-2 for 12 additional days. Tregs were stimulated using anti-CD3/anti-CD28 Dynabeads (Thermo) at a 1:1 bead:cell ratio and maintained in 300 U/mL IL-2. Following two days of culture, the stimuli were removed and cells allowed to expand in heavy or light SILAC media with 300 U/mL IL-2 for 9 days and then restimulated with more beads at 1:1 bead:cell ratio. At day 14, Tregs were counted and put into co-culture with CD8 T cells at a 1:1 Treg: CD8^+^ T cell ratio.

### Baseline CD8^+^ activation

CD8^+^ T cells were stimulated using anti-CD3/anti-CD28 Dynabeads (Thermo) at a 1:1 bead:cell ratio and maintained in 50 U/mL IL-2. Following two days of culture, the stimuli were removed and cells allowed to expand in heavy or light SILAC media with 50 U/mL IL-2 for 12 additional days, at which point CD8^+^ T cells were counted and were re-stimulated 1:1 with anti-CD3/anti-CD28 Dynabeads for three days. Isotopically-labeled activated and resting cells were then combined 1:1 for downstream processing.

### Co-culture

On day 14, isotopically-labeled CD8^+^ T cells were put into co-culture with Tregs at a 1:1 Treg:CD8^+^ T cell ratio. In addition, at the time of initiation of co-culture, anti-CD3/anti-CD28 Dynabeads were added at a 1:1 bead:CD8^+^ T cell ratio. At the time of co-culture, the CD8^+^ T cells grown in heavy SILAC media were co-cultured with Tregs, while CD8^+^ T cells were kept in monoculture and grown in light SILAC media. At the end of three days of co-culture, CD8^+^ T cells were isolated from co-culture using STEMCELL CD8^+^ enrichment kits and then combined in equal numbers with the CD8 T cells grown in mono-culture in the light SILAC media. This was done in the opposite combination of light/heavy SILAC media as well to ensure there was no bias in the experiment in assigning light or heavy SILAC media to those T cells grown in co- or mono-culture.

### Hypoxic T cell activation

To model activation in hypoxic conditions, expanded, SILAC-labeled CD8^+^ or CD4^+^ cells were collected, resuspended in fresh SILAC media, and stimulated using anti-CD3/anti-CD28 Dynabeads at a 1:10 bead:cell ratio. Cells were then either cultured at 37°C, 5% CO_2_ in normoxic (20% O_2_) or hypoxic (1% O_2_) conditions for three days. Hypoxic culture was performed in a Coy Laboratory Products hypoxic cabinet using a nitrogen/5% CO_2_ balance blend. Cells were then separated from the Dynabeads and heavy and light cells mixed at a 1:1 ratio in both forward and reverse SILAC mode before surface protein capture. For flow cytometry of IL-18R1 and CD70, cells were similarly isolated and activated but were cultured in STEMCELL Immunocult-XF T cell expansion media. Cells were stained with GHOST viability dye from Tonbo Biosciences and flow cytometry was performed using anti-IL18R1 [H44] and anti-CD70 [113-16] antibodies, both from Biolegend. Flow cytometry data was analyzed using FlowJo (v10.7.1).

### Cell surface capture

Cell surface glycoproteins were captured as previously described.^32^ Briefly, immediately after combining isotopically-labeled cells, the cells were washed in PBS, pH 6.5 and glycoproteins oxidized with 1.6 mM NaIO_4_ (Sigma) in PBS, pH 6.5 for 20 minutes at 4°C. Oxidized vicinal diols were subsequently biotinylated with 1 mM biocytin hydrazide (Biotium) in the presence of 10 mM aniline (Sigma) in PBS, pH 6.5 for 90 minutes at 4°C. Cells were then flash frozen and stored at −80°C before further preparation. To isolate glycoproteins for mass spectrometry, cell pellets were lysed with commercial RIPA buffer (VWR) supplemented with 1X Protease Inhibitor Cocktail (Sigma) and 1 mM EDTA (Sigma) for 30 minutes at 4°C. Cells were further disrupted with probe sonication and biotinylated glycoproteins pulled down with NeutrAvidin coated agarose beads (Thermo) for one hour at 4°C. Beads were transferred to Poly-Prep chromatography columns (Bio-Rad, Hercules, CA) and sequentially washed with RIPA (PBS pH 7.4 with 0.5% sodium deoxycholate [Thermo], 0.1% sodium dodecylsulfate [Fisher Scientific, Waltham, MA], 1% Nonidet P-40 substitute [VWR]), high salt PBS (PBS pH 7.4, 2 M NaCl [Sigma]), and denaturing urea buffer (50 mM ammonium bicarbonate, 2 M Urea). After washing, beads were collected and glycoproteins reduced with 5 mM TCEP (Calbiochem) for 30 minutes at 37°C and alkylated with 11 mM iodoacetamide (Sigma) for 30 minutes at room temperature. Beads were washed with urea buffer and trypsinized on-bead overnight at room temperature with 20 μg trypsin (Promega). The next day, the tryptic fraction was collected using Pierce Spin Columns before the beads were again transferred to PolyPrep columns and washed with RIPA, high salt buffer, and urea buffer before a final wash with 50 mM ammonium bicarbonate. Beads were transferred to a fresh tube and glycopeptides liberated with 5000 U/mL PNGaseF for 3 hours at 37°C. This PNGaseF fraction was collected as above. Both tryptic and PNGaseF fractions were then desalted with SOLA HRP SPE columns (Thermo) following standard protocols, dried, and dissolved in 0.1% formic acid, 2% acetonitrile prior to LC-MS/MS analysis.

### Mass spectrometry

Mass spectrometry was performed as previously described^32^, with some slight adjustments. All peptides were separated using an UltiMate 3000 UHPLC system (Thermo) with pre-packed 0.75mm x 150mm Acclaim Pepmap C18 reversed phase columns (2μm pore size, Thermo) and analyzed on a Q Exactive Plus (Thermo Fisher Scientific) mass spectrometer. For tryptic fractions, 1 μg of resuspended peptides was injected and separated using a linear gradient of 3-35% solvent B (solvent A: 0.1% formic acid, solvent B: 80% acetonitrile, 0.1% formic acid) over 230 mins at 300 μL/min. Due to the low peptide yield of the PNGase fraction, the entire fraction was injected and subsequently separated using the same gradient over 170 mins. Data-dependent acquisition was performed using a top 20 method (dynamic exclusion 35 seconds; selection of peptides with a charge of 2, 3, or 4). Full spectra with a resolution of 140,000 (at 200 m/z) were gathered in MS1 using an AGC target of 3e6, maximum injection time of 120 ms, and scan range of 400 - 1800 m/z. Centroided data from MS2 scans were collected at a resolution of 17,500 (at 200 m/z) with an AGC target of 5e4 and maximum injection time of 60 milliseconds. The normalized collision energy was set at 27 and an isolation window of 1.5 m/z with an isolation offset of 0.5 m/z was used.

### Data analysis/Statistics

SILAC proteomics data were analyzed as previously described^32^. Briefly, each individual dataset was searched for peptides using ProteinProspector v5.13.2 against the human proteome (Swiss-prot database, August 3, 2017 release). Enzyme specificity was set to trypsin with up to two missed cleavages. Cysteine carbamidomethyl was set as the only fixed modification; methionine oxidation, N-terminal glutamate to pyroglutamate, and lysine/arginine SILAC labels were set as variable modifications. Asparagine deamidation was also listed as a variable modification for the PNGaseF fractions. During the search, the peptide mass tolerance was 6 ppm, fragment ion mass tolerance was 0.4 Da, and peptide identification was filtered by peptide score of 0.0005 in ProteinProspector, resulting in a false discovery rate (FDR) of <1% calculated using the number of decoy peptides in the SwissProt database. Skyline (UWashington)^71^ software was used to perform quantitative analysis of SILAC ratios using an MS1 filtering function against a curated list of extracellular proteins generated via searches for “membrane” but not “mitochondrial” or “nuclear” using UniProt subcellular localization annotations, as previously described.^32^ For datasets collected in forward and reverse SILAC mode, spectral libraries of experiments were analyzed simultaneously to allow MS1 peaks without an explicit peptide ID to be quantified using an aligned peptide retention time. The Skyline report was subsequently exported for ratiometric analysis using a previously reported custom R script^32^. Briefly, low quality identifications (isotope dot product <0.8) were removed. For the tryptic fraction, only proteins with two or more peptides were included in downstream analysis, whereas for the PNGase fraction, only peptides with N to D deamidation were included. Ratios derived from both fractions were then combined, centered on a mean of zero, and presented as median log_2_ enrichment values. Significance was determined using a Mann-Whitney test of peptide ratios for all peptides for a given protein. Keratin 2, vimentin, and prothrombin showed dramatic enrichment in some light-labeled SILAC samples, suggesting these were contaminants and were therefore removed from downstream analysis. Heatmaps comparing expression levels between donors were generated using heatmapper.ca and other graphs were generated using GraphPad Prism (v8).

RNAseq data for naïve and activated CD8^+^ T cells was downloaded from the Database of Immune Cell eQTLs, Expression, and Epigenomics (DICE)^39^. Expression data was gathered for all overlapping proteins found in the CD8^+^ activation surfaceomics dataset, and an average expression level was calculated from all available donors in the DICE database. The expression ratio between activated and naïve was then calculated and compared with enrichments observed in the surfaceomics data. STRING analysis was performed using the online STRING Database (v11.0) and visualized using Cytoscape (v3.7.2). Gene-set enrichment analysis was performed using GSEA (v4.0.1) and the compiled data for all detected proteins within each dataset. The biological process annotated gene set (c5.bp.v7.0.entrez.gmt) used was obtained from the MSigDB collection. The CD8^+^ monoculture activation dataset was compared with both the Treg co-culture and hypoxia datasets to find pathways commonly regulated by the two immunosuppressive stimuli tested.

## Data availability

The raw proteomics data, peaklists, ProteinProspector results, and Skyline quantification results have been deposited to the ProteomeXchange Consortium via the PRIDE^72^ partner repository with the dataset identifier PXD024789. For the purposes of peer review, the data can be accessed using reviewer credentials (login; password: xz6yqlTq). Full outputs from SILAC analysis can be found in Supplemental Tables 1-4. Flow cytometry data and all other data presented is available upon reasonable request.

## REFERENCES

1. Quail, D. F. & Joyce, J. A. Microenvironmental regulation of tumor progression and metastasis. Nature Medicine 19, 1423–1437 (2013).

2. Zheng, Y. et al. Genome-wide analysis of Foxp3 target genes in developing and mature regulatory T cells. Nature 445, 936–940 (2007).

3. Togashi, Y., Shitara, K. & Nishikawa, H. Regulatory T cells in cancer immunosuppression — implications for anticancer therapy. Nat Rev Clin Oncol 16, 356–371 (2019).

4. Wing, K. et al. CTLA-4 Control over Foxp3+ Regulatory T Cell Function. Science 322, 271–275 (2008).

5. Deaglio, S. et al. Adenosine generation catalyzed by CD39 and CD73 expressed on regulatory T cells mediates immune suppression. J. Exp. Med. 204, 1257–1265 (2007).

6. Jarnicki, A. G., Lysaght, J., Todryk, S. & Mills, K. H. G. Suppression of antitumor immunity by IL-10 and TGF-beta-producing T cells infiltrating the growing tumor: influence of tumor environment on the induction of CD4+ and CD8+ regulatory T cells. J. Immunol. 177, 896–904 (2006).

7. Collison, L. W. et al. The inhibitory cytokine IL-35 contributes to regulatory T-cell function. Nature 450, 566–569 (2007).

8. Thornton, A. M. & Shevach, E. M. CD4+CD25+ Immunoregulatory T Cells Suppress Polyclonal T Cell Activation In Vitro by Inhibiting Interleukin 2 Production. Journal of Experimental Medicine 188, 287–296 (1998).

9. Grossman, W. J. et al. Human T regulatory cells can use the perforin pathway to cause autologous target cell death. Immunity 21, 589–601 (2004).

10. Sato, E. et al. Intraepithelial CD8+ tumor-infiltrating lymphocytes and a high CD8+/regulatory T cell ratio are associated with favorable prognosis in ovarian cancer. PNAS 102, 18538–18543 (2005).

11. Togashi, Y. & Nishikawa, H. Regulatory T Cells: Molecular and Cellular Basis for Immunoregulation. in Emerging Concepts Targeting Immune Checkpoints in Cancer and Autoimmunity (ed. Yoshimura, A.) 3–27 (Springer International Publishing, 2017). doi:10.1007/82_2017_58.

12. Vaupel, P. & Mayer, A. Hypoxia in Tumors: Pathogenesis-Related Classification, Characterization of Hypoxia Subtypes, and Associated Biological and Clinical Implications. in Oxygen Transport to Tissue XXXVI (eds. Swartz, H. M., Harrison, D. K. & Bruley, D. F.) 19–24 (Springer, 2014). doi:10.1007/978-1-4939-0620-8_3.

13. Muz, B., de la Puente, P., Azab, F. & Azab, A. K. The role of hypoxia in cancer progression, angiogenesis, metastasis, and resistance to therapy. Hypoxia (Auckl) 3, 83–92 (2015).

14. Madsen, C. D. et al. Hypoxia and loss of PHD2 inactivate stromal fibroblasts to decrease tumour stiffness and metastasis. EMBO Rep. 16, 1394–1408 (2015).

15. Branco-Price, C., Evans, C. E. & Johnson, R. S. Endothelial hypoxic metabolism in carcinogenesis and dissemination: HIF-A isoforms are a NO metastatic phenomenon. Oncotarget 4, 2567–2576 (2013).

16. Eales, K. L., Hollinshead, K. E. R. & Tennant, D. A. Hypoxia and metabolic adaptation of cancer cells. Oncogenesis 5, e190–e190 (2016).

17. Hu, K. H. et al. ZipSeq: barcoding for real-time mapping of single cell transcriptomes. Nat Methods 17, 833–843 (2020).

18. Lewis, C. & Murdoch, C. Macrophage Responses to Hypoxia: Implications for Tumor Progression and Anti-Cancer Therapies. The American Journal of Pathology 167, 627–635 (2005).

19. Zhang, Y. & Ertl, H. C. J. Starved and Asphyxiated: How Can CD8(+) T Cells within a Tumor Microenvironment Prevent Tumor Progression. Front Immunol 7, 32 (2016).

20. Dang, E. V. et al. Control of T(H)17/T(reg) balance by hypoxia-inducible factor 1. Cell 146, 772–784 (2011).

21. Westendorf, A. M. et al. Hypoxia Enhances Immunosuppression by Inhibiting CD4+ Effector T Cell Function and Promoting Treg Activity. Cell. Physiol. Biochem. 41, 1271–1284 (2017).

22. Vuillefroy de Silly, R., Dietrich, P.-Y. & Walker, P. R. Hypoxia and antitumor CD8+ T cells: An incompatible alliance? Oncoimmunology 5, (2016).

23. Hatfield, S. M. et al. Systemic oxygenation weakens the hypoxia and hypoxia inducible factor 1α-dependent and extracellular adenosine-mediated tumor protection. J Mol Med 92, 1283–1292 (2014).

24. Fischer, K. et al. Inhibitory effect of tumor cell–derived lactic acid on human T cells. Blood 109, 3812–3819 (2007).

25. Hatfield, S. M. et al. Immunological mechanisms of the antitumor effects of supplemental oxygenation. Science Translational Medicine 7, 277ra30–277ra30 (2015).

26. Scharping, N. E. et al. Mitochondrial stress induced by continuous stimulation under hypoxia rapidly drives T cell exhaustion. Nat Immunol 22, 205–215 (2021).

27. Rodriguez-Garcia, A., Palazon, A., Noguera-Ortega, E., Powell, D. J. J. & Guedan, S. CAR-T Cells Hit the Tumor Microenvironment: Strategies to Overcome Tumor Escape. Front. Immunol. 11, (2020).

28. Ahmadzadeh, M. et al. Tumor antigen–specific CD8 T cells infiltrating the tumor express high levels of PD-1 and are functionally impaired. Blood 114, 1537–1544 (2009).

29. Ribas, A. & Wolchok, J. D. Cancer immunotherapy using checkpoint blockade. Science 359, 1350–1355 (2018).

30. Wollscheid, B. et al. Mass-spectrometric identification and relative quantification of N-linked cell surface glycoproteins. Nature Biotechnology 27, 378–386 (2009).

31. Martinko, A. J. et al. Targeting RAS-driven human cancer cells with antibodies to upregulated and essential cell-surface proteins. eLife 7, e31098 (2018).

32. Leung, K. K. et al. Multiomics of azacitidine-treated AML cells reveals variable and convergent targets that remodel the cell-surface proteome. PNAS 116, 695–700 (2019).

33. Pfistershammer, K. et al. CD63 as an Activation-Linked T Cell Costimulatory Element. The Journal of Immunology 173, 6000–6008 (2004).

34. Bayer, A. L., Baliga, P. & Woodward, J. E. Transferrin receptor in T cell activation and transplantation. J. Leukoc. Biol. 64, 19–24 (1998).

35. Vomhof-DeKrey, E. E., Haring, J. S. & Dorsam, G. P. Vasoactive Intestinal Peptide Receptor 1 is Downregulated During Expansion of Antigen-Specific CD8 T Cells Following Primary and Secondary Listeria monocytogenes Infections. J Neuroimmunol 234, 40–48 (2011).

36. Spendlove, I. & Sutavani, R. The role of CD97 in regulating adaptive T-cell responses. Adv. Exp. Med. Biol. 706, 138–148 (2010).

37. Lin, M. et al. Cell surface antigen CD109 is a novel member of the α2 macroglobulin/C3, C4, C5 family of thioester-containing proteins. Blood 99, 1683–1691 (2002).

38. Ren, W. et al. Amino-acid transporters in T-cell activation and differentiation. Cell Death Dis 8, e2655–e2655 (2017).

39. Schmiedel, B. J. et al. Impact of Genetic Polymorphisms on Human Immune Cell Gene Expression. Cell 175, 1701–1715.e16 (2018).

40. Ivetic, A., Hoskins Green, H. L. & Hart, S. J. L-selectin: A Major Regulator of Leukocyte Adhesion, Migration and Signaling. Front. Immunol. 10, (2019).

41. Lauer, V. et al. Hypoxia drives glucose transporter 3 expression through HIF-mediated induction of the long non-coding RNA NICI. J. Biol. Chem. jbc.RA119.009827 (2019) doi:10.1074/jbc.RA119.009827.

42. Wang, H. et al. Cellular Hypoxia Promotes Heterotopic Ossification by Amplifying BMP Signaling. J Bone Miner Res 31, 1652–1665 (2016).

43. Liberzon, A. et al. The Molecular Signatures Database Hallmark Gene Set Collection. cels 1, 417–425 (2015).

44. Munkley, J. & Elliott, D. J. Hallmarks of glycosylation in cancer. Oncotarget 7, 35478–35489 (2016).

45. Grandi, A. et al. ERMP1, a novel potential oncogene involved in UPR and oxidative stress defense, is highly expressed in human cancer. Oncotarget 7, 63596–63610 (2016).

46. Bensellam, M. et al. Hypoxia reduces ER-to-Golgi protein trafficking and increases cell death by inhibiting the adaptive unfolded protein response in mouse beta cells. Diabetologia 59, 1492–1502 (2016).

47. Ye, J. et al. ER stress induces cleavage of membrane-bound ATF6 by the same proteases that process SREBPs. Mol. Cell 6, 1355–1364 (2000).

48. Abdul Rahim, S. A. et al. Regulation of hypoxia-induced autophagy in glioblastoma involves ATG9A. Br. J. Cancer 117, 813–825 (2017).

49. Renna, M. et al. Autophagic substrate clearance requires activity of the syntaxin-5 SNARE complex. J Cell Sci 124, 469–482 (2011).

50. O’Neill, R. E. et al. T Cell-Derived CD70 Delivers an Immune Checkpoint Function in Inflammatory T Cell Responses. J. Immunol. 199, 3700–3710 (2017).

51. Manieri, N. A., Chiang, E. Y. & Grogan, J. L. TIGIT: A Key Inhibitor of the Cancer Immunity Cycle. Trends in Immunology 38, 20–28 (2017).

52. Deng, J. et al. Hypoxia-induced VISTA promotes the suppressive function of myeloid-derived suppressor cells in the tumor microenvironment. Cancer Immunol Res (2019) doi:10.1158/2326-6066.CIR-18-0507.

53. O’Neill, R. E. et al. T Cell–Derived CD70 Delivers an Immune Checkpoint Function in Inflammatory T Cell Responses. The Journal of Immunology 199, 3700–3710 (2017).

54. Wang, B. et al. Combination cancer immunotherapy targeting PD-1 and GITR can rescue CD8+ T cell dysfunction and maintain memory phenotype. Science Immunology 3, (2018).

55. Raczkowski, F. et al. CD39 is upregulated during activation of mouse and human T cells and attenuates the immune response to Listeria monocytogenes. PLOS ONE 13, e0197151 (2018).

56. Martin, M. et al. CD84 Functions as a Homophilic Adhesion Molecule and Enhances IFN-γ Secretion: Adhesion Is Mediated by Ig-Like Domain 1. The Journal of Immunology 167, 3668–3676 (2001).

57. Kim, M. N. et al. Activated Leukocyte Cell Adhesion Molecule Stimulates the T-Cell Response in Allergic Asthma. Am. J. Respir. Crit. Care Med. 197, 994–1008 (2018).

58. Qualai, J. et al. Expression of CD11c Is Associated with Unconventional Activated T Cell Subsets with High Migratory Potential. PLOS ONE 11, e0154253 (2016).

59. Bettelli, E. et al. Integrin alpha 4 differentially affect the migration of effector and regulatory T cells (P4113). The Journal of Immunology 190, 133.10–133.10 (2013).

60. Wei, C.-W. et al. Equilibrative Nucleoside Transporter 3 Regulates T Cell Homeostasis by Coordinating Lysosomal Function with Nucleoside Availability. Cell Reports 23, 2330–2341 (2018).

61. Macintyre, A. N. et al. The Glucose Transporter Glut1 is Selectively Essential for CD4 T Cell Activation and Effector Function. Cell Metab 20, 61–72 (2014).

62. Doedens, A. L. et al. Hypoxia-inducible factors enhance the effector responses of CD8^+^ T cells to persistent antigen. Nature Immunology 14, 1173–1182 (2013).

63. Gropper, Y. et al. Culturing CTLs under Hypoxic Conditions Enhances Their Cytolysis and Improves Their Anti-tumor Function. Cell Reports 20, 2547–2555 (2017).

64. Choi, S.-H. et al. Hypoxia-induced RelA/p65 derepresses SLC16A3 (MCT4) by downregulating ZBTB7A. Biochimica et Biophysica Acta (BBA) - Gene Regulatory Mechanisms 1862, 771–785 (2019).

65. Watson, M. J. et al. Metabolic support of tumour-infiltrating regulatory T cells by lactic acid. Nature 1–7 (2021) doi:10.1038/s41586-020-03045-2.

66. Renner, K. et al. Restricting Glycolysis Preserves T Cell Effector Functions and Augments Checkpoint Therapy. Cell Reports 29, 135–150.e9 (2019).

67. Mazure, N. M. & Pouysségur, J. Hypoxia-induced autophagy: cell death or cell survival? Current Opinion in Cell Biology 22, 177–180 (2010).

68. Pino, S. C. et al. Protein kinase C signaling during T cell activation induces the endoplasmic reticulum stress response. Cell Stress Chaperones 13, 421–434 (2008).

69. MacIver, N. J., Michalek, R. D. & Rathmell, J. C. Metabolic Regulation of T Lymphocytes. Annual Review of Immunology 31, 259–283 (2013).

70. Roth, T. L. et al. Reprogramming human T cell function and specificity with non-viral genome targeting. Nature 559, 405–409 (2018).

71. Pino, L. K. et al. The Skyline ecosystem: Informatics for quantitative mass spectrometry proteomics. Mass Spectrom Rev 39, 229–244 (2020).

72. Perez-Riverol, Y. et al. The PRIDE database and related tools and resources in 2019: improving support for quantification data. Nucleic Acids Res 47, D442–D450 (2019).

