## Supplemental Materials for "Hypoxia is a dominant remodeler of the CD8^+^ T cell surface proteome relative to activation and regulatory T cell-mediated suppression"

### SUPPLEMENTAL FIGURES

**Figure S1. Expansion of T cells in SILAC media effectively labels proteins.**

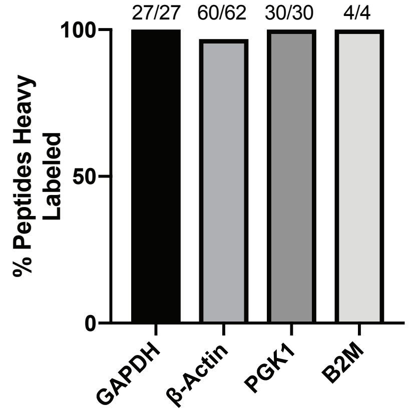


**Supplemental Figure 1. Expansion of T cells in SILAC media effectively labels proteins.** Representative data from one CD8$^+$ T cell labeling experiment following 2 weeks of expansion in heavy SILAC media. Cells were lysed, proteins trypsinized, peptides desalted, and analyzed with LC-MS/MS. Numbers above bars indicate the number of heavy labeled peptides identified for each protein out of the total number of peptides for that protein.

Figure S2. CD8+ activation in monoculture donor comparisons.

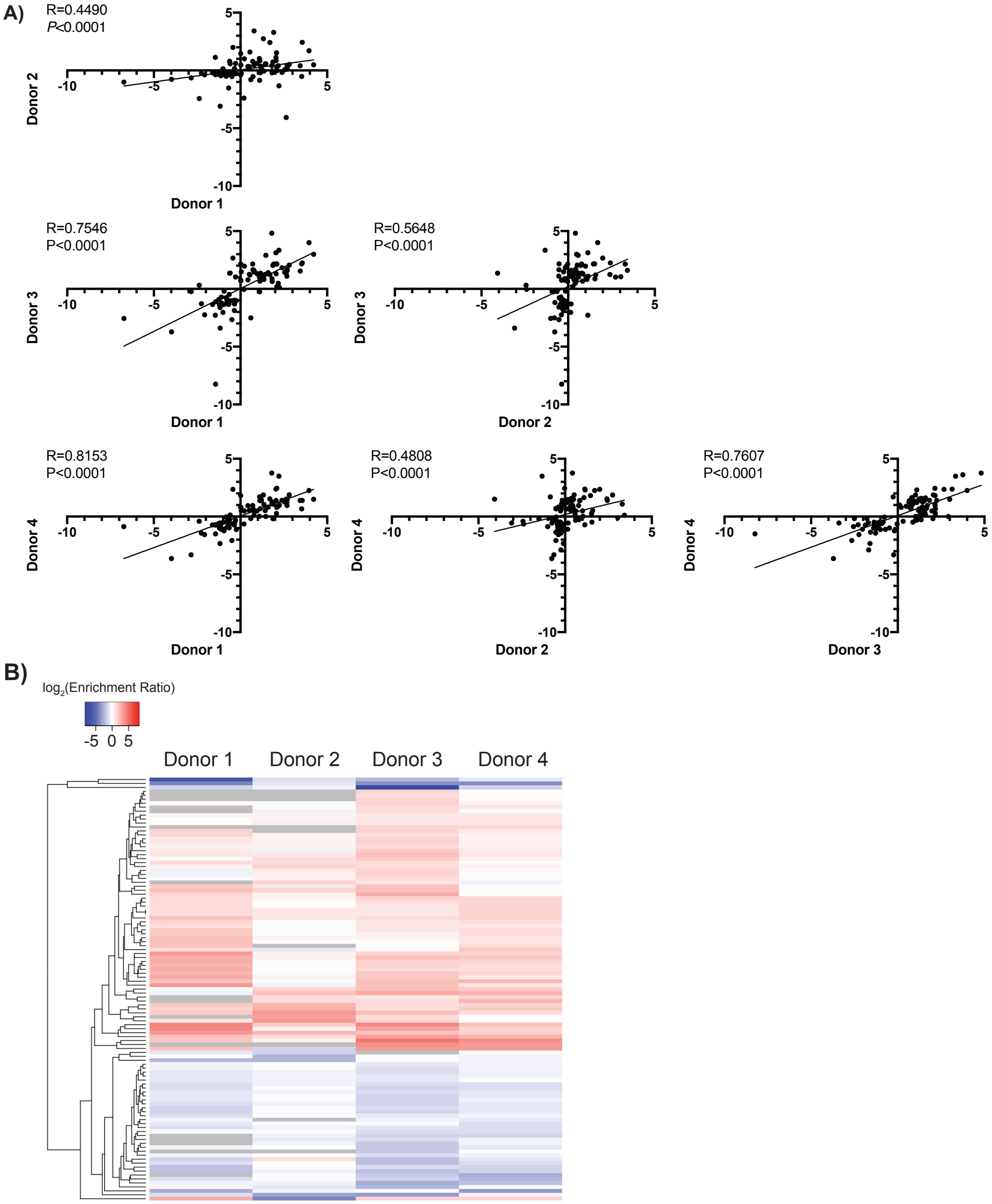


**Supplemental Figure 2. CD8$^+$ activation in monoculture donor comparisons.** (A) Spearman correlations comparing SILAC ratios of all significantly-altered proteins identified when analyzing compiled data from N=4 donors. (B) Heatmap showing SILAC ratio for significantly-altered proteins for each donor. Grey boxes indicate the protein was not identified in cells from that donor.

**Figure S3. Correlation of $CD8^+$ activation proteomics data with RNAseq data from the DICE Database.**

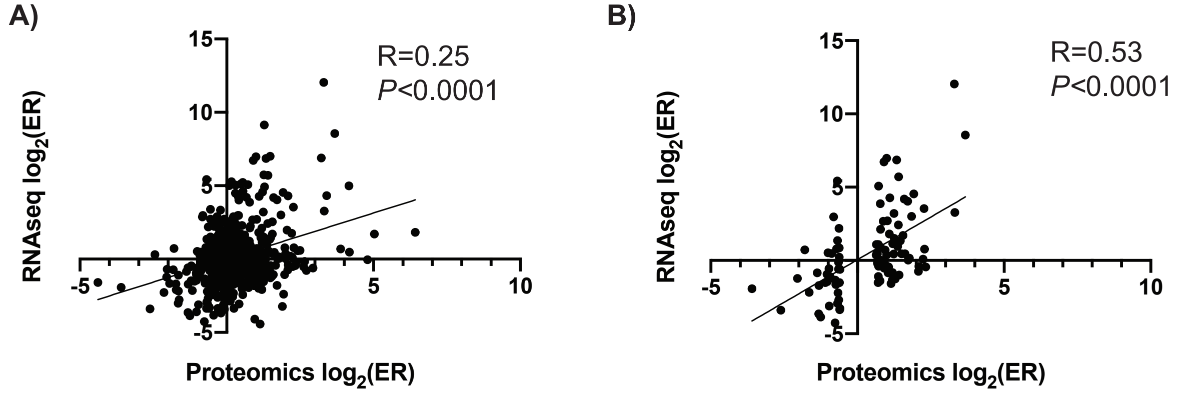


**Supplemental Figure 3. Correlation of $CD8^+$ activation proteomics data with RNAseq data from the DICE Database.** Correlations for all proteins (A) and significantly-altered proteins (B) from the activation proteomics dataset with activation data from the DICE Database. Expression data from the DICE Database was averaged for all replicates, then a $log_2$(enrichment ratio [ER]) was calculated by dividing the expression signal for activated naïve $CD8^+$ cells by the signal for resting naïve $CD8^+$ cells. Only proteins found in both datasets are shown.

**Figure S4. CD8⁺ activation in Treg co-culture donor comparisons.**

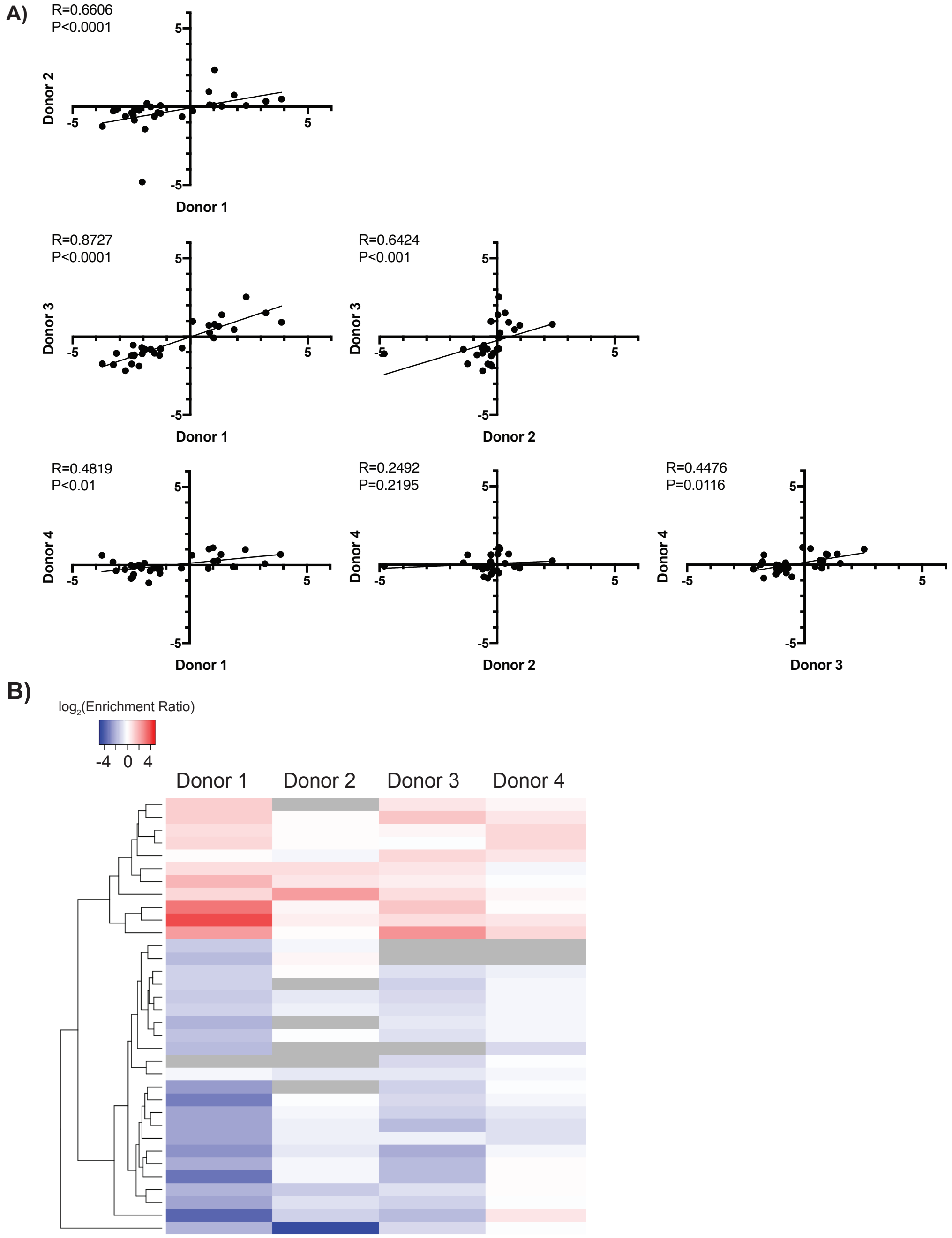


**Supplemental Figure 4. CD8$^+$ activation in Treg co-culture donor comparisons.** (A) Spearman correlations comparing SILAC ratios of all significantly-altered proteins identified when analyzing compiled data from N=4 donors. (B) Heatmap showing SILAC ratio for significantly-altered proteins for each donor. Grey boxes indicate the protein was not identified in cells from that donor.

Figure S5. Effect of hypoxia on CD8⁺ T cell expansion and viability.

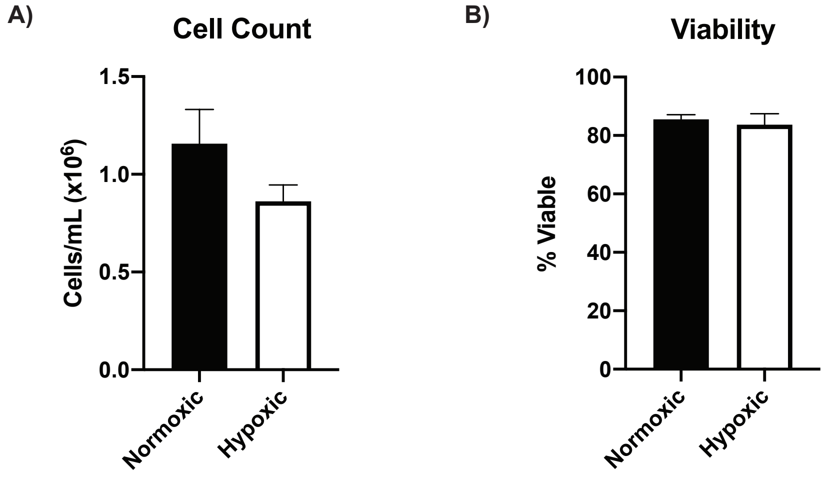


**Supplemental Figure 5. Effect of hypoxia on $CD8^+$ T cell expansion and viability.** Bar graphs showing cell counts (A) and viability (B) of $CD8^+$ cultures after three days of activation in either normoxic or hypoxic conditions. Data represent mean -/+ standard error of the mean for N=3 biological replicates, each with two technical replicates.

**Figure S6. CD8⁺ activation in hypoxia donor comparisons.**

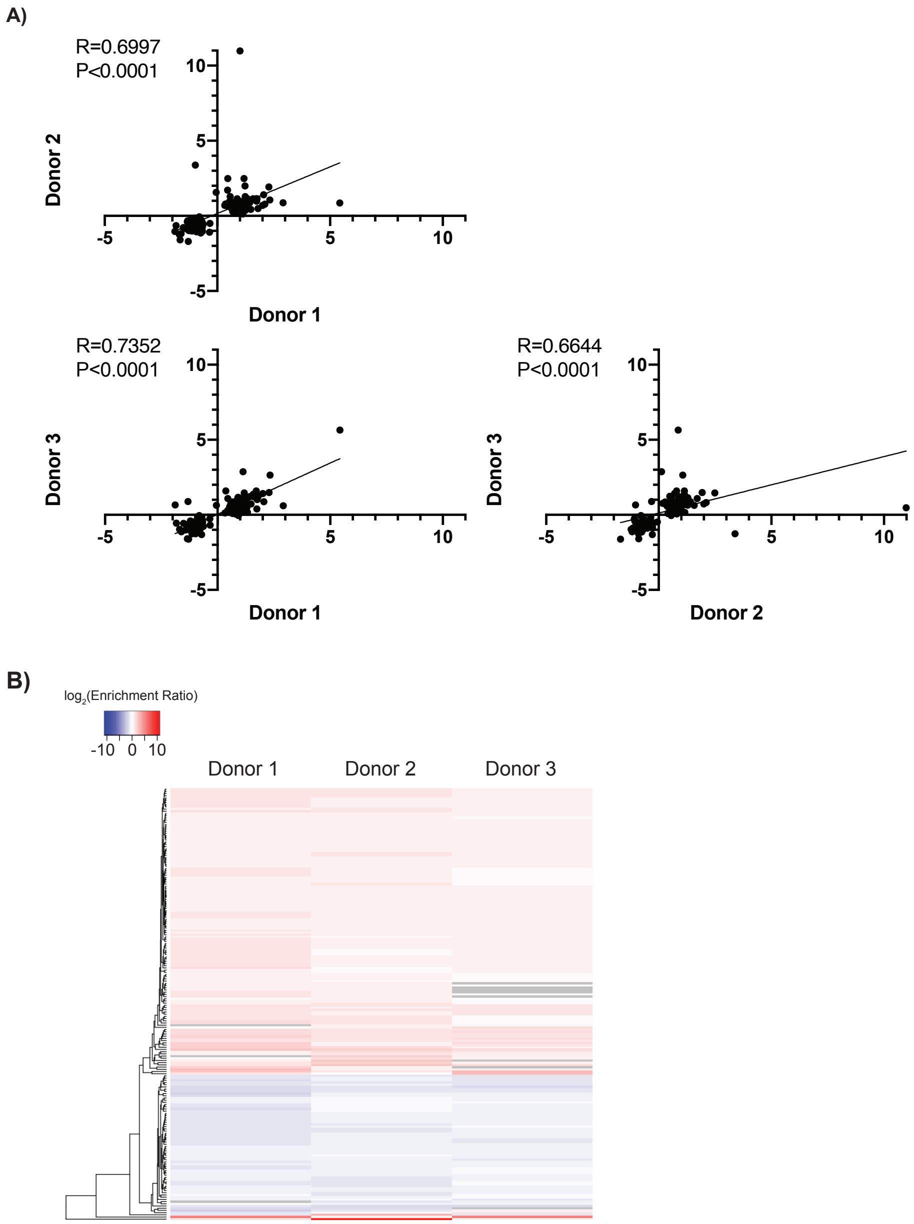


**Supplemental Figure 6. CD8$^+$ activation in hypoxia donor comparisons.** (A) Spearman correlations comparing SILAC ratios of all significantly-altered proteins identified when analyzing compiled data from N=3 donors. (B) Heatmap showing SILAC ratio for significantly-altered proteins for each donor. Grey boxes indicate the protein was not identified in cells from that donor.

Figure S7. Flow cytometry of IL18R1 and CD70 on resting, activated, and hypoxic $CD8^+$ T cells.

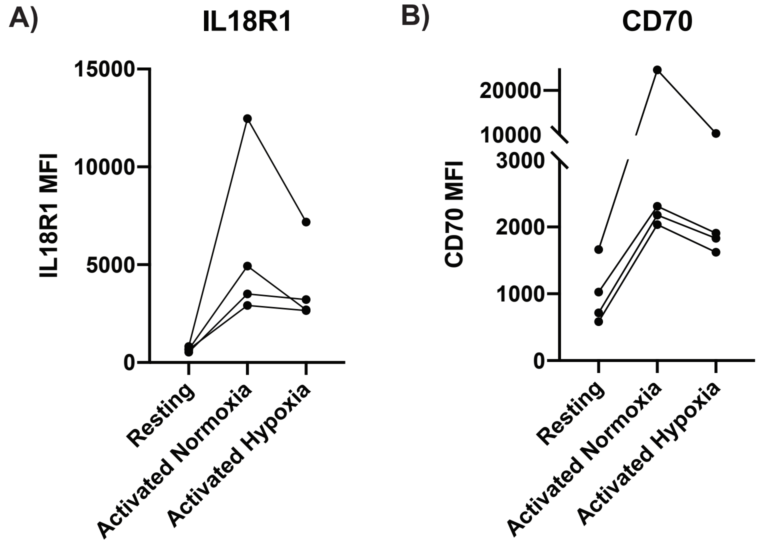


**Supplemental Figure 7. Flow cytometric analysis of IL18R1 and CD70 expression on hypoxic $CD8^+$ T cells.** Cells from N=4 donors were either left resting or stimulated in normoxic (20% $O_2$) or hypoxic (1% $O_2$) for three days. Cells were then collected and stained for either IL18R1 (A) or CD70 (B). Mean fluorescence intensity (MFI) values are shown for each donor. Lines connect datapoints from the same donor.

### SUPPLEMENTAL DATA FILE LEGENDS

**Supplemental Table 1. SILAC analysis output of $CD8^+$ activation in monoculture experiments.** Excel sheet contains tabs with the raw output of our in-house analysis script showing SILAC enrichment ratio and *P*-value for identified proteins. Additional tabs show significantly ($P$<0.05, -/+ 1.5-fold change) up- and downregulated protein lists.

**Supplemental Table 2. SILAC analysis output of $CD8^+$ activation in Treg co-culture experiments.** Excel sheet contains tabs with the raw output of our in-house analysis script showing SILAC enrichment ratio and *P*-value for identified proteins. Additional tabs show significantly ($P$<0.05, -/+ 1.5-fold change) up- and downregulated protein lists.

**Supplemental Table 3. SILAC analysis output of $CD8^+$ activation in hypoxia experiments.** Excel sheet contains tabs with the raw output of our in-house analysis script showing SILAC enrichment ratio and *P*-value for identified proteins. Additional tabs show significantly ($P$<0.05, -/+ 1.5-fold change) up- and downregulated protein lists.

**Supplemental Table 4. SILAC analysis output of $CD4^+$ Teff activation in hypoxia experiments.** Excel sheet contains tabs with the raw output of our in-house analysis script showing SILAC enrichment ratio and *P*-value for identified proteins. Additional tabs show significantly ($P$<0.05, -/+ 1.5-fold change) up- and downregulated protein lists.
